# Tissue-resident NK cells support survival in pancreatic cancer through promotion of cDC1-CD8T activity

**DOI:** 10.1101/2023.10.14.562332

**Authors:** Simei Go, Constantinos Demetriou, Giampiero Valenzano, Sophie Hughes, Simone Lanfredini, Helen Ferry, Edward Arbe-Barnes, Shivan Sivakumar, Rachael Bashford-Rogers, Mark R. Middleton, Somnath Mukherjee, Jennifer Morton, Keaton Jones, Eric O’Neill

## Abstract

The immunosuppressive microenvironment in pancreatic ductal adenocarcinoma (PDAC) prevents tumor control and strategies to restore anti-cancer immunity (i.e. by increasing CD8 T cell activity) have had limited success. Here we demonstrate how inducing localized physical damage using ionizing radiation (IR) unmasks the benefit of immunotherapy by increasing tissue-resident NK (trNK) cells that support CD8 T activity. Our data confirms that targeting mouse orthotopic PDAC tumors with IR together with CCR5 inhibition and PD1 blockade reduces E-cadherin positive tumor cells by recruiting a hypoactive NKG2D^-ve^ NK population, phenotypically reminiscent of trNK cells, that supports CD8 T cell involvement. We show an equivalent population in human PDAC cohorts that represents immunomodulatory trNK cells that could similarly support CD8 T cell levels in a cDC1-dependent manner. Importantly, a trNK signature associates with survival in PDAC and solid malignancies revealing a potential beneficial role for trNK in improving adaptive anti-tumor responses and supporting CCR5i/αPD1 and IR-induced damage as a novel therapeutic approach.

## Introduction

Over the past decade, immune checkpoint inhibitors have shown significant success in the treatment of various solid malignancies. Treatment of pancreatic cancer using immunotherapy alone or in combination with radiotherapy or chemotherapy has unfortunately seen limited clinical success over either modality alone (Bockorny et al., 2020; Doi et al., 2019; Royal et al., 2010; Weiss et al., 2018; Weiss et al., 2017) (Mohindra et al., 2015). This is likely due to several tumor microenvironmental factors, including the dense stroma that supports an immunosuppressive environment while also obstructing the infiltration of cytotoxic immune cells (Mills et al., 2022; Piper et al., 2023). Novel strategies comprising dual targeting of (programmed cell death-1) PD-1/IL-2Rβγ with radiotherapy are beginning to indicate that potential benefits may require a coordinated alteration in the suppressive microenvironment involving loss of regulatory T cells (Tregs) and increased natural killer (NK) cell infiltration, in addition to simply increasing CD8 T cell infiltration (Piper et al., 2023).

We previously identified serum CCL5 (RANTES) as a negative prognostic marker for late-stage advanced pancreatic cancer (Willenbrock et al., 2021). CCL5 is pro-inflammatory and an acute response to injury or infection that promotes recruitment of monocytes and lymphoid immune cells through the receptors CCR5, CCR3, and CCR1. In cancer, sustained CCL5-mediated inflammation leads to a suppressive environment via attraction of CCR5^+^ immunoregulatory components including Tregs, tumor-associated macrophages (TAMs), and myeloid-derived suppressor cells (MDSCs) normally involved in resolving inflammatory events (Hemmatazad & Berger, 2021). Pancreatic tumors that are poorly differentiated produce higher levels of CCL5 and express more of its main receptor, CCR5, compared to well-differentiated and non-cancerous tissue (Monti et al., 2004; Singh et al., 2018). Disrupting this loop by the FDA-approved CCR5 inhibitor maraviroc reduces pancreatic tumor cell migration, invasiveness (Singh et al., 2018) and proliferation (Huang et al., 2020). Silencing CCL5 expression in Panc02 cells or systemic administration of the CCR5 inhibitor TAK-779 reduces Treg migration and tumor volume in a murine subcutaneous tumor model (Tan et al., 2009). These studies suggest that the CCL5/CCR5 axis may be a promising therapeutic target in pancreatic cancer as it has the potential to alter both the intrinsic properties of tumor cells and immune cell migration.

In established pancreatic tumors, the immune suppressive environment promotes more tissue repair/resolution signaling and prevents pro-inflammatory signaling (Mantovani et al., 2017). Radiotherapy is delivered to localized tumors to stimulate a pro-inflammatory environment and increase the opportunity for tumor neo-antigens to be recognized by infiltrating adaptive response cells (Benkhaled et al., 2023). As such, stereotactic ablative radiotherapy (SBRT) delivers high doses of radiotherapy in a minimal number of fractions to maximize an inflammatory cascade but the sensitivity of lymphocytes to DNA damage and the immune suppressive environment prevent benefit or meaningful tumor control(Mills et al., 2022). In NSCLC, single cell RNA sequencing (scRNA-seq) of immune cells within tumors identifies a distinct subset of CD49a^+^ CD103^+^ tissue-resident memory (TRM) CD8^+^ T cells that have capacity to respond to neo-antigens but are functionally suppressed (Caushi et al., 2021). In this case, the use of an anti-PD1 antibody (αPD1) supports TRM activation but additional disruption of the immune suppressive environment is still required, indicating that additional components are required in addition to simply activating CD8^+^ T cell neo-antigen recognition.

Kirchhammer et al. recently reported that a tissue-resident NK (trNK) cell population is induced in NSCLC in response to viral delivery of IL-12, which crucially supported type I conventional dendritic cell (cDC1) infiltration and increased DC-CD8^+^ T cell interactions (Kirchhammer et al., 2022). Together with PD1 blockade, IL-12-mediated recruitment of trNK cells enhances cross-presentation of antigen to CD8 T via cDC1, suggesting this represents a substantial barrier for T cell focused therapy and that improving the NK/DC/T-cell crosstalk can promote antitumor immunity and tumor control. Notably, this was dependent on CCL5 and points to a positive role for inflammatory CCL5 signaling in the absence of CCR5-mediated suppression (Kirchhammer et al., 2022).

Given the negative prognosis associated with high CCL5 serum levels in human patients and its diverse functions in chemokine and paracrine signaling (Aldinucci et al., 2020), we hypothesized that inhibiting CCR5 rather than CCL5, in combination with radiotherapy and anti-PD1 immunotherapy may improve tumor control via a multimodal approach. To test this, we employed a murine orthotopic pancreatic cancer model and monitored tumor growth and immune infiltration. We find that while the use of a CCR5 inhibitor (CCR5i) alone restricts Treg involvement, it does not therapeutically beneficially impact tumor viability alone or together with αPD1. However, in combination with radiotherapy, we see a significant alteration in MDSCs, NK and CD8 T cells and better tumor control. Interestingly, we observe that IR/CCR5i/αPD1 combination treatment induced a trNK population in which the NKG2D^-^ NK cells are the highly correlated immune population with tumor control. Exploration of single-cell RNA sequencing (scRNA-seq) datasets from human PDAC studies confirms the presence of trNK cells as immunomodulatory in human PDAC, and in which these cells seem to directly support cDC1-CD8 communication. Strikingly, a specific trNK signature indicates that higher levels of this NK subtype are significantly correlated with better PDAC patient overall and disease-free survival. Moreover, pan-cancer analysis reveals that trNK cell involvement is associated with better patient survival across a number of solid tumors and supports the potential utility of a combination regimen comprising ionizing radiation (IR) and CCR5i/αPD1 immunotherapy (IT) as a promising strategy to increase NK/cDC1/CD8 mediated tumor control in solid cancers.

## Results

### CCL5 is a negative prognostic marker in pancreatic cancer

We previously identified serum CCL5 as a *bona fide* negative prognostic marker for pancreatic cancer(Willenbrock et al., 2021), and found that, in two independent pancreatic ductal adenocarcioma (PDAC) cohorts (CPTAC3 (Cao et al., 2021) and TCGA (Uhlen et al., 2017)), higher CCL5 expression associates with poor overall and disease-free survival, confirming the negative implication of elevated levels of CCL5 in pancreatic cancer (Figure 1A). To understand the cause of CCL5-mediated reduced survival in PDAC, we hypothesized that immune cells responsive to a CCL5 chemotactic gradient (through expression of the cognate receptors CCR5, CCR3 or CCR1) could be potential contributors to an adverse tumor immune environment (Figure 1B, Figure supplement 1A). Single cell RNA sequencing of human tumors has revealed genetic signatures for myeloid derived suppressor cells of polymorphonuclear (PMN-MDSC) or monocytic (M-MDSC) origin (Alshetaiwi et al., 2020), TAMs (Wang et al., 2021), CD4 regulatory T cells (Treg)(Mijnheer et al., 2021) and NK cells (Smith et al., 2020), but none of these immune populations have yet been implicated as a causative agent in the poor outcome associated with high CCL5 expression in PDAC (Figure 1C). Notably, exploration of individual genes within these signatures indicated stark opposing correlations within each signature pool – particularly for genes associated with MDSCs (S100P vs ARG2) and NK cells (CD56 vs CD16) (Figure 1C, Figure supplement 1B). CD56 (*NCAM1*) is a key marker that represents functionally distinct subpopulations of NK cells where CD56^bright^ NK cells represent immature states that move to CD56^dim^ upon maturation to full cytotoxic potential, whereas conversely CD16^+^ expression marks activation and CD16^-^ immature or quiescent NK cells (Poli et al., 2009). Thus, these distinct subtypes may differentially contribute to survival, e.g., association of CD16 with poor survival may implicate a detrimental role of CD56^dim^CD16^+^ NK cells, whereas the strong positive prognosis associated with CD56 expression could indicate benefit of CD56^bright^CD16^-^ NK cells. The benefit of CD56 can be attributed to NK cells over neuronal expression as the neuronal-cell specific homolog *NCAM2* has no prognostic value (Figure supplement 1C).

**Figure 1.**
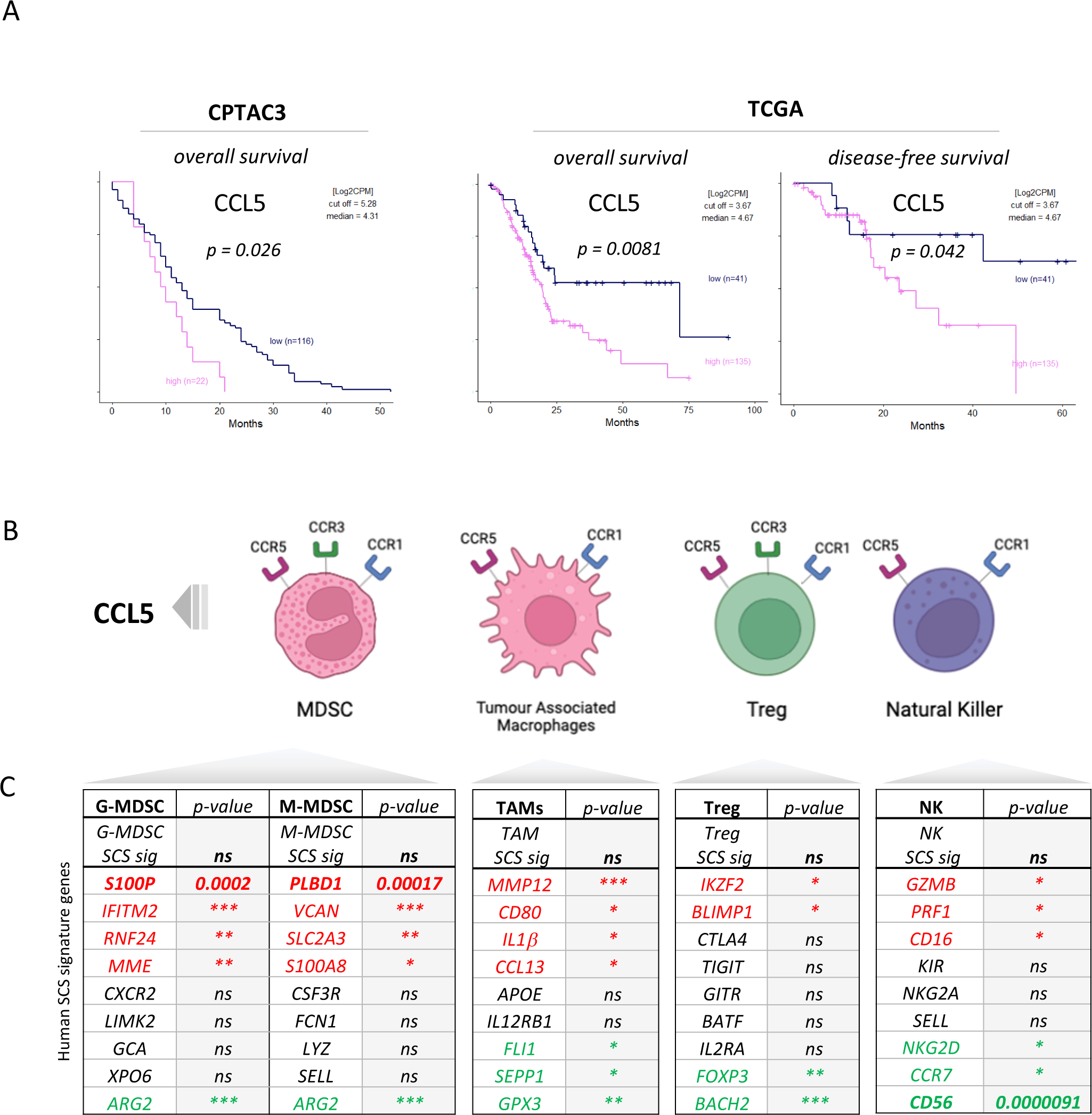
(A) Overall and disease-free Kaplan-Meier survival plots of PDAC patients segregated into high or low CCL5 gene expression levels within pancreatic tumors. Data are derived from CTPAC3 and TCGA cohorts and optimal cut-off values were calculated using the max-stat method for each respective cohort. (B) Schematic overview of CCL5-responsive immune cells and corresponding CCL5 receptor repertoire expression. (C) Correlation between overall and single cell gene signatures of CCL5-responsive immune cells with overall PDAC prognosis. Colour depicts positive (green), negative (red) or neutral (black) prognostic outcomes (*p<0.05, ** p<0.01, *** p<0.005). Data are derived from the Pathology Dataset of the Human Protein Atlas and based on human tissue micro arrays and correlated log-rank p-value for Kaplan Meier analysis

### CCR5i modulates Treg infiltration in a murine orthotopic pancreatic tumor model

We next employed a syngeneic orthotopic pancreatic cancer model where cells derived from *Kras^G12D^*, *Trp53^R172H^*, *Pdx1-Cre* (KPC) tumor bearing mice are injected into the pancreas of wildtype C57BL/6 mice, thereby recapitulating a human pancreatic cancer microenvironment (Matzke-Ogi et al., 2016). From a selection of independently isolated KPC derived cell lines, we first determined the cell line KPC-F as an appropriate model for human PDAC tumor based on expression of epithelial (E-cadherin), mesenchymal (vimentin), and stromal (Collagen I, αSMA) markers as well as displaying growth kinetics (determined by MRI) amenable for the study and expression of high levels of CCL5 (Figure 2A, Figure supplement 2A/B). To address which immune subsets are involved in CCL5 signaling in pancreatic cancer, we developed a 17-color spectral flow cytometry panel to monitor the tumor-infiltrating immune microenvironment in parallel (Figure supplement 2C) and employed a specific CCR5 inhibitor (maraviroc, CCR5i), to block CCR5, the most widespread and highest-affinity receptor for CCL5 expressed on Tregs (Figure supplement 1A).

**Figure 2.**
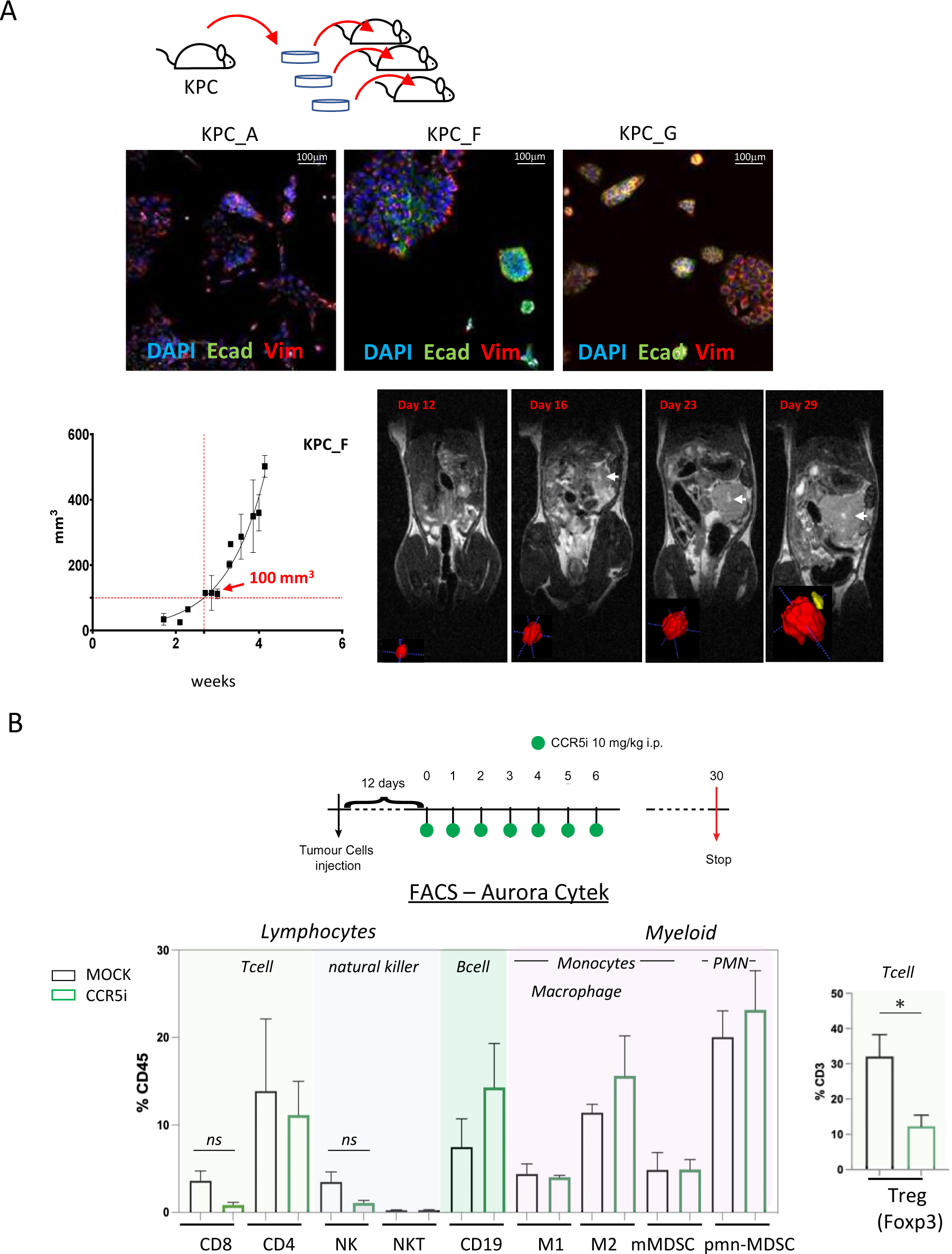
(A) Three different lineages of KPC pancreatic tumor cells (derived from Kras^G12D^p53^R172H^Pdx1-Cre mice) were obtained and stained for DAPI (blue; nucleus), E-cadherin (green; epithelial) and Vimentin (red; mesenchymal). Growth curve of orthotopically injected KPC-F cells (500 cells) into the pancreas of wildtype C57BL/6 over time in weeks. Tumor volume was measured using MRI. Representative MRI images over time are displayed, white arrow denotes tumor mass. (B) Timeline of maraviroc (CCR5i) treatment regimen. A total of 12 days post-orthotopic injection, tumor-bearing mice were treated daily with maraviroc (10 mg/kg via intraperitoneal injection) for 6 days and followed for up to 30 days after starting treatment. Frequencies of pancreatic tumor-infiltrating immune cells harvested at day 30 with or without maraviroc using spectral flow cytometry is shown. Data are represented as mean percentage positive cells of Live/CD45^+^cells±SD. For Tregs, the mean percentage positive cells of Live/CD45^+^ CD3^+^ cells± SD is shown. Significance was tested using the Welch and Brown-Forsythe ANOVA for parametric data or Kruskal-Wallis test for non-parametric data. Mock (n=6), IR (n=3), aPD1 (n=8), aPD1+IR (n=8), CCR5i (n=3), CCR5i+IR (n=8), aPD1+CCR5i (n=5), αPD1/CCR5i/IR (n=8).

Pilot experiments injecting 500 KPC-F cells yielded robust and replicative growth kinetics (indicated by matching volumes of ±100 mm^3^ at the beginning of exponential growth), permitting direct comparisons of infiltrating immune cells across different treatment groups (Figure 2A). Next, tumors were allowed to reach 50 mm^3^ (approx. 12 days) before initiation of treatment with daily administration of CCR5i for 7 days. Mice were culled 30 days post-treatment for the characterization of infiltrating immune cells (Figure 2B). Mice in the CCR5i-treated group showed a trend towards reduction in NK cells and all T cell subsets with a significant reduction in Treg (Live/CD45^+^CD3^+^CD4^+^FOXP3^+^) infiltration, indicating the active recruitment of CCR5^+^ Tregs in PDAC (Figure 2B). Given the enhanced effect of CCR5i on tumor growth kinetics (Figure 3A) however, standalone inhibition of CCR5 is unlikely to clinically benefit PDAC progression.

**Figure 3.**
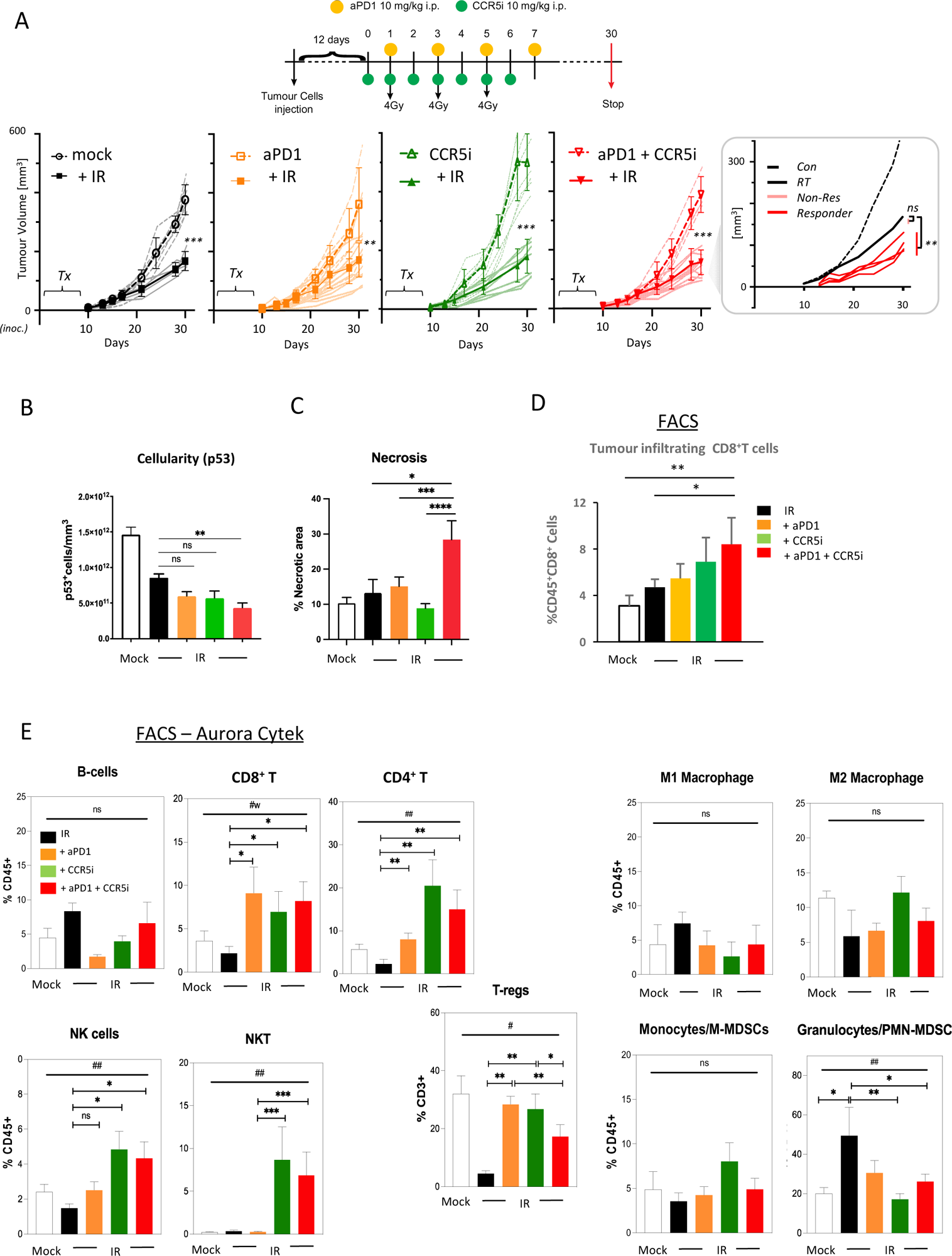
(A) Timeline of triple treatment regimen (maraviroc, αPD1 and radiotherapy) following orthotopic injection of KPC-F cells. A total of 12 days post orthotopic injection of 500 KPC-F cells in the pancreas of wildtype C57BL/6 mice, mice were treated as follows: seven consecutive days of 10 mg/kg intraperitoneal injection of maraviroc and four alternating days of 10 mg/kg intraperitoneal injection of αPD1. Mice were followed for up to 30 days following the start of the treatment regimen. Tumor volumes were measured by MRI and growth curves of individual treatment groups are plotted with or without radiotherapy as measured by MRI. Average growth curves±SD are depicted in bold, individual mice are shaded (without IR; dashed, with IR; solid). Insert: expanded view of triple combination to show ‘responders’ display a significant benefit over RT alone. (B) Quantification of pancreatic tumors derived from (A) stained by IHC for p53. (C) Quantification of necrotic areas in pancreatic tumors derived from (A) based on H&E staining. (D) Quantification of infiltrating CD8 T cells in pancreatic tumors derived from (A) by flow cytometry. (E) Profiling of infiltrating immune cells in pancreatic tumors derived from (A) by flow cytometry as in Figure 2B. Single, live cells were included for analysis and are represented as frequencies of Live/CD45^+^ cells or total CD3^+^ for FoxP3^+^ Tregs. Significance was tested using the Welch and Brown-Forsythe ANOVA for parametric data # p<0.05, ##p<0.01 or Kruskal-Wallis test for non-parametric data #w p<0.01; pairwise comparisons (student t test) *p<0.05, **p<0.01, ***p<0.005, ****p<0.001.

### Combination immunotherapy following localized damage alters tumor immune composition

Induction of a localized inflammatory microenvironment with ionising radiation (IR) can drive anti-tumor responses, but there is limited evidence for clinical benefit of radiotherapy in combination with immune checkpoint inhibitors in PDAC (ClinicalTrials.gov identifier: NCT04098432) (Chen et al., 2022). Therefore, we next examined whether the addition of CCR5i to tumor-targeted IR could improve this by further modulating immune cell migration, in particular by reducing Treg involvement (Figure 3A, Figure supplement 3A). As expected, radiotherapy (3 x 4Gy) produced a strong effect on gross tumor volume, significantly reducing volumes over standalone treatment with αPD1, CCR5i or CCR5i+αPD1 (Figure 3A). Responses to combination of IR with CCR5i, αPD1 or CCR5i+αPD1 (IR+IT) showed larger variations compared to IR alone, implying mixtures of ‘responders’ and ‘non-responders’ to immune-modulatory treatments. Moreover, IR/CCR5i/αPD1 treated tumor sections had significantly reduced DAPI^+^ and p53^+^ KPC cells compared to all other conditions, suggesting significantly more loss of tumor cells by triple combination treatment (Figure 3B, Figure supplement 3B, C). Tumor sections from the triple combination treatment also presented with increased loss of active stroma (αSMA staining, Figure supplement 3D) and increased necrotic areas over standalone radiotherapy (Figure 3C). In line with apparent increased tumor control, the triple combination treatment demonstrated infiltration of CD8^+^ T cells (Figure 3D), supporting greater penetration of CD8^+^ T cells into the centre of tumors (Figure supplement 3E). These results support improved tumor control with combination of IR/CCR5i/αPD1 over standalone radiotherapy or IR in combination with αPD1 and CCR5i alone. To elucidate the immune mechanisms behind this control, we analysed the immune infiltrate of pancreatic tumors. In line with the results above, the triple combination reduced Treg infiltration, enhanced CD8^+^ T cell infiltration and supported a moderate increase in CD4^+^ T helper cells (Figure 3E, Figure supplement 3F).

Interestingly, no alteration was seen in the myeloid compartment, except for a reduced infiltration of PMN-MDSCs, whereas a significant infiltration of NK and NKT cells could be observed in pairwise comparisons between IR+αPD1 vs IR+CCR5i or IR/CCR5i/αPD1 (Figure 3E, Figure supplement 3F). These results collectively support the notion that IR/CCR5i/αPD1 combination treatment alters immune infiltration by reducing Tregs and increasing NK and CD8 T cells, thereby resulting in greater local tumor control.

### CD8 T and NKG2D^-^ NK cells correlate with increased tumor control

To derive more granularity on the potential roles of NK- and T-cell populations in tumor control, tissue sections were stained using an optimized multiplex immunofluorescence panel and analyzed by HALO AI software to identify tumor cells (E-cadherin^+^ - blue), CD8^+^ T cells (CD3^+^CD8^+^ - orange), CD4^+^ T-cells (CD3^+^CD8^-^ - green) and NK cells (CD3^-^CD8^-^NK1.1^+^ - red) (Figure 4A, Figure supplement 4A). As observed with flow cytometric and immunohistochemical analyses, multiplex staining of tumor sections also revealed a significant increase of CD8^+^ T cells per mm^3^ when IR was used in combination with CCR5i or CCR5i+αPD1 (Figure supplement 4B). Notably, despite having responders and non-responders in the combination groups (Figure 4A), a significant overall reduction in E-cadherin^+^ tumor cells as a percentage of total DAPI^+^ cells was observed (Figure supplement CB), supporting a decrease in cellularity and an increase in tumor necrosis with the IR/CCR5i/αPD1 combination (Figure 3B, C). The increase in CD8^+^ T cells with combination treatment appears independent of tumor area and is matched by a similar increase in NK cells (Figure supplement 4B, C). To correlate immune infiltration against loss of tumor cells (a measure of local tumor control) we determined relationships between CD4 T, CD8 T and NK cell populations and E-cadherin^+^ cells across all tumor sections (independent of treatment) and found a significant inverse correlation for both CD8 T and NK cells (r^2^=-0.3, p=0.038 and r^2^=-0.33, p=0.026 respectively), but not CD4 T cells (Figure 4B).

**Figure 4.**
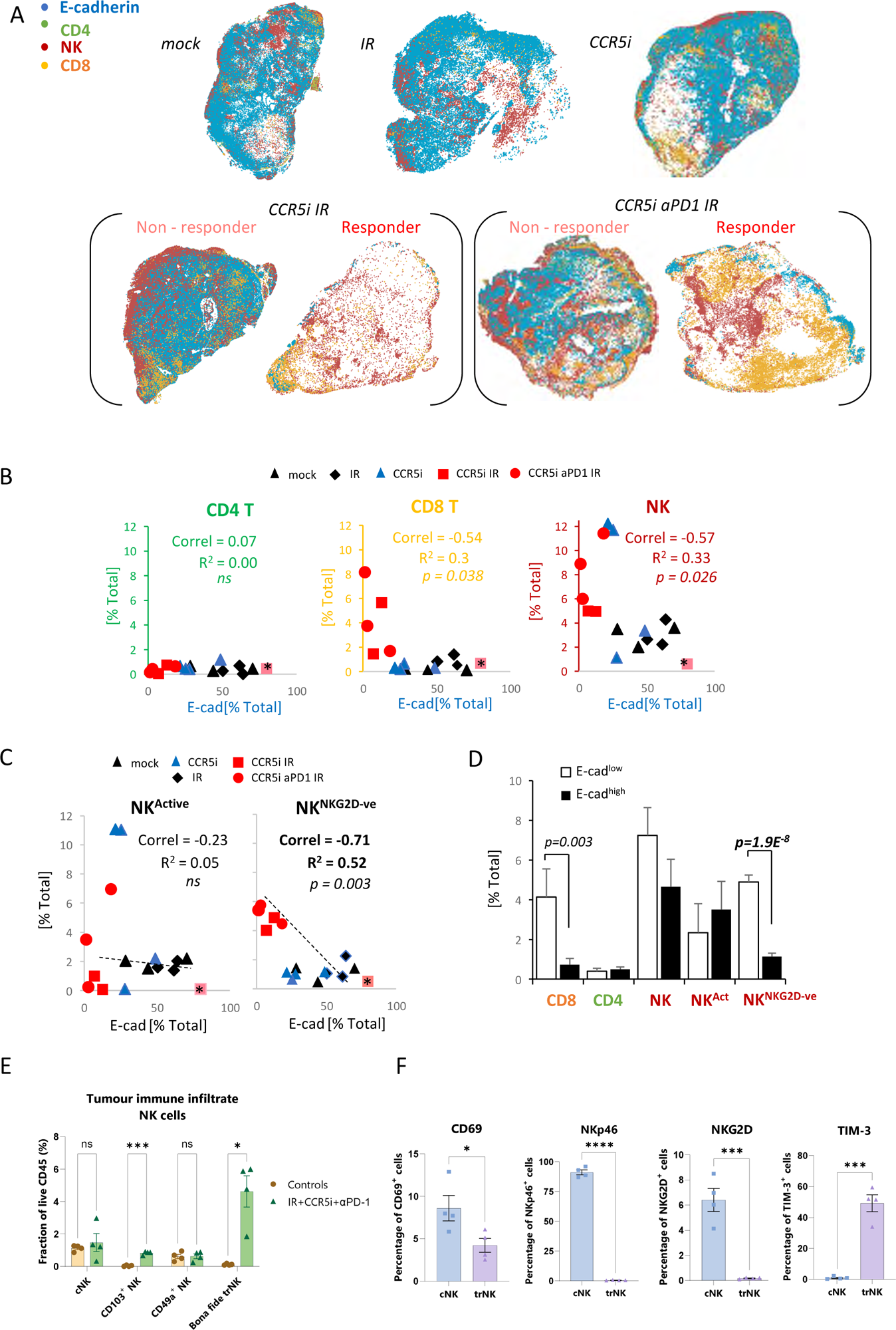
(A) Spatial plots of individual cells identified using HALO software of scanned multiplex immunofluorescence murine orthotopic pancreatic tumor slices. Positive staining is identified as the marker of interest and DAPI+ (nucleus stain) signal. Responders and non-responders to treatment are based on loss of E-cadherin staining. (B) Correlations of total CD4 T (CD3^+^CD8^-^), CD8 T (CD3^+^CD8^+^) and NK cells (CD3^-^NK1.1^+^) plotted against %positive E-cadherin^+^ cells as derived from (A). (C) Correlation of %positive segregated NK cells plotted against %positive E-cadherin+ cells as derived from (A). NK cells were segregated based on expression of NKG2D; NK^Active^; NK1.1^+^NKG2D^+^; NK^NKG2D-ve^; NK1.1^+^NKG2D^-^. (D) Intra-tumoral immune cells of stratified pancreatic tumors based on low or high E-cadherin percentage (cut-off: 20%). Significance was tested for p<0.05 with a two-tailed student’s T-test. * censored non-responder. (E) Proportion of infiltrating trNK (Live/CD45^+^CD3^-^CD19^-^NK1.1^+^CD103^+^CD49a^+^), conventional NK cells (Live/CD45^+^CD3^-^CD19^-^NK1.1^+^CD103^-^CD49a^-^), CD103^+^ NK, and CD49a^+^ NK cells isolated from orthotopic pancreatic tumors of mice treated with the IR+IT regimen and controls, as a percentage of CD45^+^ cells. Significance was tested for p<0.05 with a student’s t-test. (F) Comparative surface expression of activation marker (CD69), activating receptors (NKp46, NKG2D) and exhaustion marker (TIM-3) on cNK cells and trNK cells isolated from orthotopic pancreatic tumours. Significance was tested for p<0.05 with a student’s t-test.

We next wondered whether the association between NK cells and tumor control was due to cytotoxicity by functionally active NK cells by focusing our analysis on the NKG2D^+^ NK cell population, given that this is one of the major activating receptors on NK cells and is required for the lysis of target cells (Bryceson & Ljunggren, 2008). Surprisingly, the proportion of infiltrating CD3^-^NK1.1^+^NKG2D^+^ cells (NK^Active^) in stained tissue sections was reduced under IR/CCR5iαPD1 combination treatment, implying a decrease in total NK cell cytotoxic capacity (Figure supplement 4D). The reduced NKG2D expression on NK cells may be a result of the prolonged engagement by ligands expressed on tumor cells, followed by ligand-induced endocytosis and degradation (Quatrini et al., 2015), or the shedding of NKG2D ligands by tumor cells (Kaiser et al., 2007). In both instances, receptor down-regulation causes a reduction in cytotoxicity and impairs NK cell responsiveness to tumor cells, potentially contributing to exhaustion (Groh et al., 2002). Moreover, circulating NK cells from PDAC patients show reduced NKG2D levels compared to healthy controls (Peng et al., 2013) supporting the notion that chronic exposure to NKG2D ligands expressed or shed by PDAC cells might cause NKG2D down-modulation and a hypo-responsive phenotype. Indeed, the reciprocal CD3^-^ NK1.1^+^NKG2D^-^ population increases upon triple combination treatment (Figure supplement 4C). Surprisingly, the correlation of NK cells with decreased frequency of E-cadherin^+^ cells was completely abrogated with selection for more functionally competent NK cells (NK^Active^), implying that the opposite hypoactive NKG2D^-^ NK population (NK^NKG2D-ve^) is responsible for correlation of NK cells with E-cadherin loss and indeed, a superior inverse correlation is observed for NK^NKG2D-ve^ and E-cadherin compared to NK^Active^ or NK^Total^ (r^2^=0.52, p=0.003) (Figure 4C). To explore this association is relation to tumor control, sections were split into either E-cadherin^high^ or E-cadherin^low^ and the extent of immune involvement was represented as a percentage of total DAPI^+^. While CD8 T cells significantly segregated with E-cadherin^low^, implying a contribution to tumor control, the association of NK^NKG2D-ve^ was vastly more significant (Figure 4D).

To further dissect the association between NK cells and loss of tumor cells, we immunophenotyped NK cells infiltrating tumors of mice treated with the IR/CCR5i/αPD1 regimen with an NK-cell dedicated spectral flow cytometry panel. Using a combination of Uniform Manifold Approximation and Projection (UMAP) dimensionality reduction and manual gating, we found that more than two thirds of tumor-infiltrating NK cells expressed the canonical markers of tissue-residency CD103 and CD49a (Figure supplement 4E, F), with the double positive population (hereafter referred to as *bona fide* tissue resident NK cells, or trNK) being barely detectable in untreated mice and rising nearly 40-fold after treatment (Figure 4E). The remainder cells expressed neither marker of tissue-residency and were, therefore, named conventional NK cells (or cNK).

We then compared these two clusters and found that trNK were less likely to be activated and expressed significantly lower levels of the activating receptors NKG2D and NKp46, as well as significantly higher levels of the inhibitory receptor TIM-3 (Figure 4F). In line with our multiplex IF data (Figure supplement 4D), these data indicate that NK cells infiltrating tumors of mice treated with the IR/CCR5i/αPD1 combination were predominantly tissue resident and hypoactive, with a minority displaying a more conventional, fully active phenotype.

### Heterogenous subsets of NK cells in PDAC exhibit differential inhibitory and activating signatures

To extrapolate our findings to the human setting, we explored scRNA-seq data from isolated NK and T-cells derived from human pancreatic cancer patients. A total of 51,561 cells were catalogued into 17 distinct cell lineages annotated with canonical gene signatures as described in Steele *et al*. using unsupervised clustering (Steele et al., 2020) (Figure supplement 5A). UMAP projection of the lymphocyte compartment did not delineate NK subpopulations and showed overlap with CD8 T cells (Figure supplement 5B). Therefore, we focused on UMAP projections of the CD8 and NK compartment alone, which led to three clear NK subpopulations that are distinct from CD8 T cells and clearly separate out in clusters of hypoactive (NK_C1), fully active (NK_C2), and cytotoxic (NK_C3) NK cells from T cells (Figure 5A). In agreement with downregulation of circulating NKG2D^+^ in PDAC patients, NKG2D (KLRK1) expression was below detection across all CD8 and NK subpopulations retrieved from patient tumors(Groh et al., 2002). As expected, CD16^high^ (FCGR3a) cytolytic NK cells (NK_C3) cluster separately from CD16^-^ NK cells (NK_C1) which are more enriched for genes related to cytokine secretion than cytolytic function (Figure 5D, Figure supplement 5C). The NK_C1 cluster correlates best with the tissue-resident, hypoactive NK phenotype observed in mice as they similarly displayed reduced cytolytic (reduced NKG7, NKp80, GZMA and PRF1) with additional expression of tissue residency markers *CD103*, *CD49a* and the adaptive activating receptor NKG2C (*KLRC2*) (Figure 5B, C). While adaptive NK cells in the peripheral circulation are associated with a CD56^dim^CD16^+^ phenotype, adaptive-like (i.e. NKG2C^+^) CD56^bright^CD16^dim^ trNK cells have been identified in other epithelial sites such as the lungs (Brownlie, Scharenberg, Mold, Hård, et al., 2021), therefore, our pancreatic tissue NK_C1 cells could comprise, at least in part, adaptive-like tissue-resident NK cells. Downstream UMAP analysis of the NK cell population distinguishes these 3 subsets of NK cells (NK_C1, NK_C2 and NK_C3) based on 41 differentially expressed genes (Figure 5D, E).

**Figure 5.**
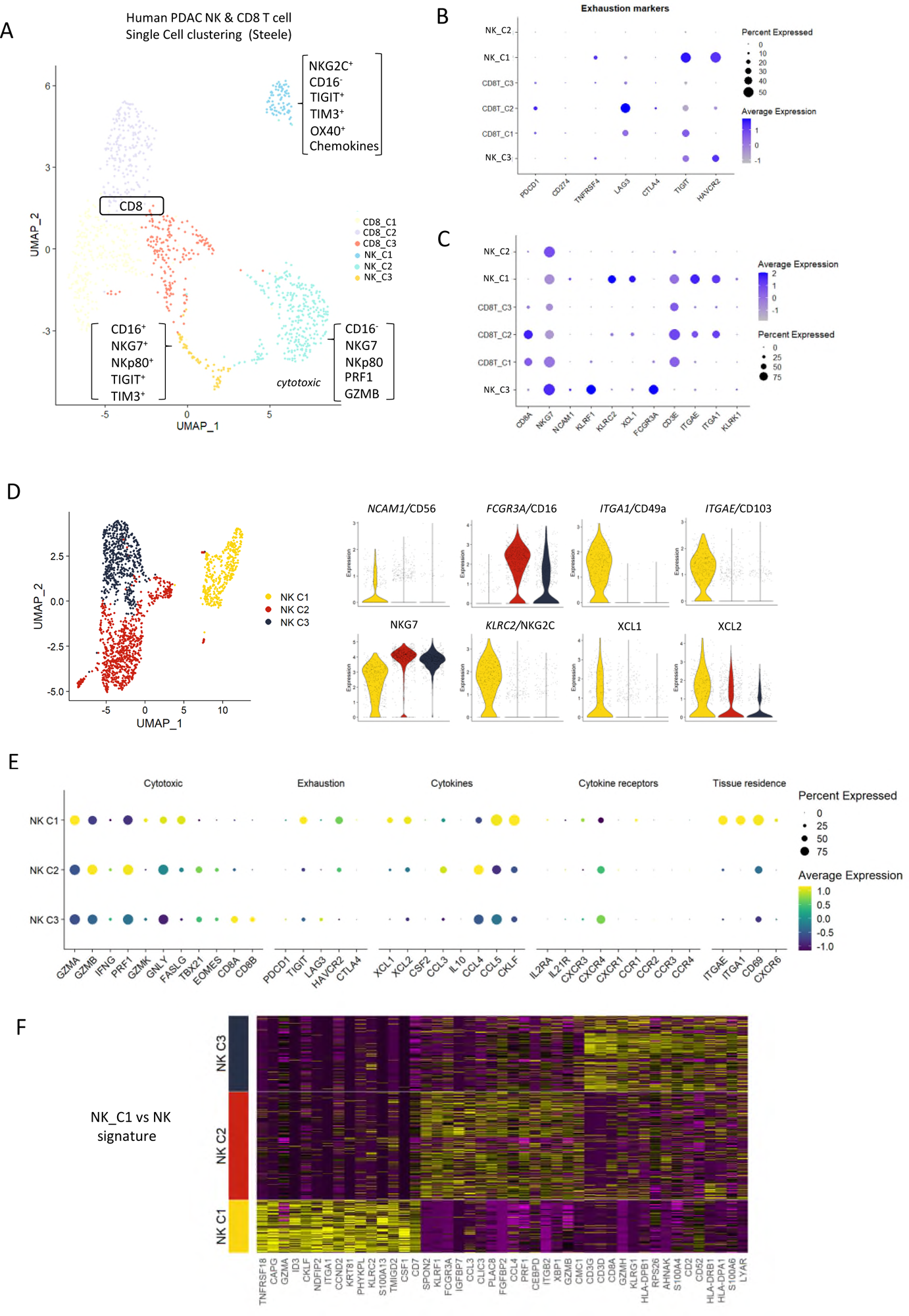
(A) UMAP of the CD8+ T and NK sub-clusters from Steele et al. (B) Dot plot showing the expression of exhaustion-related genes across CD8+ T and NK cell sub-clusters. (C) Dot plot showing highly expressed genes for each sub-cluster. (D) UMAP of the three NK subclusters (left), and violin plots comparing the expression of NK subtype-associated genes between the sub-clusters (right). (E) Dot plot showing the different gene expression programs across the three NK sub-clusters. (F) Heatmap showing the top 15 upregulated markers for each NK sub-cluster compared against total NK cluster.

Given that CD56 expression correlates with increased survival in PDAC patients (Figure 1C) we were intrigued to notice that the hypoactive NK_C1 cluster is enriched for CD56 but not CD16 expression (Figure 5D). These data suggest that the NK_C1 cluster represents a subset of NK cells in PDAC that display tissue-residency markers (*ITGAE,* CD103 and *ITGA1,* CD49a) and have an immune-secretory phenotype (XCL1) as opposed to cytotoxic phenotype (NK_C2 CD16^+^, NKG7^+^) (Figure 5E). Furthermore, we defined a core signature gene set to distinguish NK_C1 from other NK subpopulations (Figure 5F). Verification of the NK_C1 population as trNK cells in a second scRNA-seq pancreatic cancer dataset of Peng *et al*. (Peng et al., 2013) further supports the existence of these hypoactive cells in an independent PDAC cohort (Figure supplement 5D, E, F).

### Tissue resident NK cells in PDAC show differential communication

The NK_C1 population had the highest expression of the chemokines XCL1 and XCL2, which have been demonstrated to attract type-1 conventional dendritic cells (cDC1) (Böttcher et al., 2018) and thereby increase cross presentation to CD8^+^ T cells. We next explored myeloid populations in the Steele *et al*. scPDAC dataset (Figure supplement 6A) and identified cDC1 cells as XCR1^+^ enriched compared to other DC subsets (Figure 6A, Figure supplement 6B). As XCR1 is the receptor for XCL1/2, this suggests that one of the main directions of communication from NK_C1 cells in PDAC is to cDC1 (Figure 6B). We next employed the R-package ‘CellChat’ (Jin et al., 2021) to further dissect the crosstalk between NK_C1 and other cells found in the pancreatic tumor microenvironment. We again observed that NK_C1-derived XCL1/2 is the strongest signal to XCR1^+^ cDC1s (Figure 6C), supporting active crosstalk between NK_C1 and cDC1. Analysis of specific ligand-receptor pairs between NK_C1 and 24 other cell groups yielded 32 significant interactions, of which the TNFSF14 (LIGHT)-TNFRSF14 pair was the most universal intercellular interaction, whereas CD74 (or MIF) signaling showed the highest communication probability (Figure supplement 6C).

**Figure 6.**
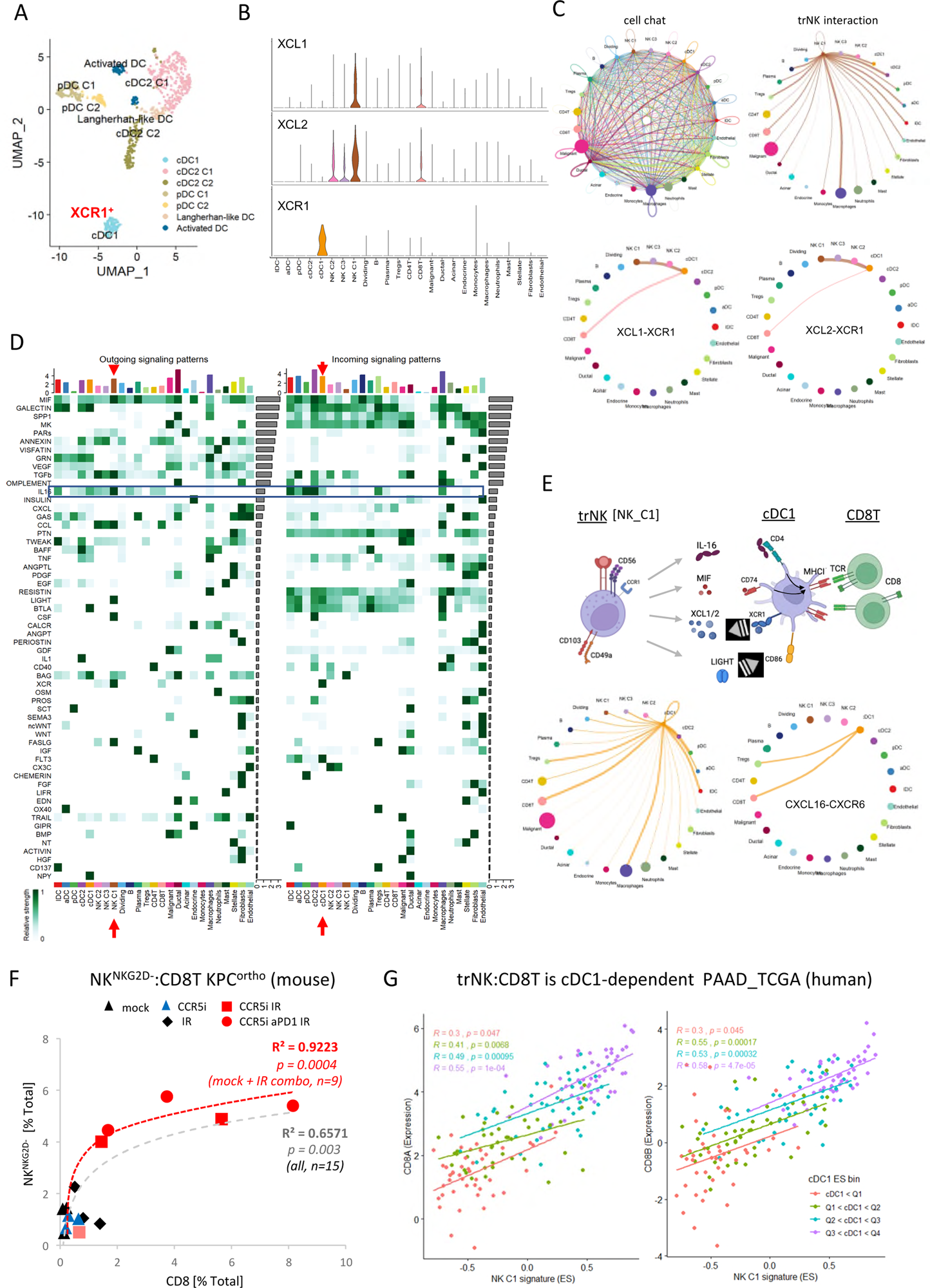
(A) UMAP of the dendritic cell sub-clusters from the Steele dataset. (B) Violin plot showing the expression of XCL1, XCL2, and XCR1 across all cell types. (C) Circle plots showing interactions across all cell types (top left), signals coming from the tissue-resident NK cells (top right), the XCL1-XCR1 interaction (bottom left), and the XCL2-XCR1 interaction (bottom right). The width of edges represents the communication strength. (D) Heatmap showing the summary of secreted signalling communications of outgoing (left) and incoming (right) signals. The colour bar represents the communication strength, and the size of the bars represents the sum of signalling strength for each pathway or cell type. (E) Schematic overview of the trNK to cDC1 and cDC1 to CD8 T cell communication axis (top). Circle plots of all outgoing signals from cDC1 (bottom left) and the CXCL16-CXCR6 signalling (bottom right). (F) Correlation of HALO data on total NK^NKG2D-ve^ NK cells (CD3^-^NK1.1^+^NKG2D^-^) with CD8 T cells (CD3^+^CD8^+^) from stained sections of treated KPC_F orthotopic tumors, R^2^ and p-values indicate positive correlation across all tumors (gray, n=15) or limited to mock, CCR5i+IR and IR/CCR5i/αPD1 combination (red, n =9). (G) Correlation of trNK signature with CD8A (left) or CD8B (right) in bulk RNA-seq from TCGA_PAAD and binned into quartiles based on extent of cDC1 involvement as assumed by cDC1 signature (Table supplement 1).

Within the immune cell compartment, communication signals between NK_C1 and macrophages were the most abundant and diverse, followed by cDCs (cDC2 and cDC1) and Tregs (Figure supplement 6C). Similarly, cDC1 in these PDAC samples also seem capable of presenting MHC class I peptides to CD8^+^ T cells, but surprisingly also MHC class II to CD4^+^ Tregs (Figure supplement 6D). Next, significant receptor-ligand interactions were segregated as ‘outgoing’ or ‘incoming’ signals to understand directionality of communication. Among the outgoing signals from different cell types, NK_C1 cells contributed the highest IL-16, CSF, LIGHT (TNFSF14), FASLG and MIF signals in tumors (Figure 6D). IL-16 is mainly known as a chemoattractant for CD4^+^ T cells, however CD4^+^ dendritic cells have also been demonstrated to be recruited by IL-16 (Bialecki et al., 2011; Kaser et al., 1999; Vremec et al., 2000). IL-16 has also been described to increase HLA-DR levels in CD4^+^ T cells and eosinophils, indicating that it has the potential to induce MHC antigen presentation in CD4^+^ cells (Cruikshank et al., 1987; Rand et al., 1991). Taken together, these interactions support the hypothesis that trNK cells may improve tumor control via recruitment of cDC1 via XCR1, while promoting DC maturation via LIGHT-CD86 signaling (Zou & Hu, 2005) and supporting DC antigen presentation to both CD4^+^ and CD8^+^ T cells via MIF-CD74 signaling (Basha et al., 2012) (Figure 6E). On the other hand, dendritic cell-secreted BAG6 could promote both survival and cytokine release of NK cells by binding to NKp30 (Simhadri et al., 2008) and directly signal to CD8 T cells via CXCL16-CXCR6 (Di Pilato et al., 2021; Vella et al., 2021), thereby generating an anti-tumor feedforward loop (Figure 6E). IL-16 also communicates to CD4^+^ Tregs, which could contribute to immunosuppression via enhanced Treg migration and expansion (Figure 6D)(McFadden et al., 2007). However, Tregs may also be susceptible to Fas-mediated cell death due to NK_C1 expression of Fas ligand (Figure 6D). This could enhance immune cell infiltration and revoke the immunosuppressive environment, ultimately contributing to increased tumor control.

These results so far suggest the presence of immunoregulatory trNK cells in PDAC that are involved in an intricate immune communication network with DCs and CD8 T cells to enhance anti-tumor immunity. To support this hypothesis, we explored the correlation between trNK and CD8 T infiltration in our murine orthotopic tumor model and found a highly significant positive association (R² = 0.6571, p = 0.003) which was strengthened when we focused on untreated (Mock) versus IR+IT combinations where trNK cells are evident (R² = 0.9223, p = 0.0004) (Figure 6F). To ascertain if this also holds true in human PDAC, we first specified the NK_C1 signature as a 14-gene signature that was specific for our trNK cells over all other cells to interrogate these cells in bulk datasets (Table supplement 1). We next explored the CD8 T:trNK cell relationship in PAAD_TCGA and, as the model in Figure 6E predicts this interaction to be cDC1 dependent, binned the PAAD_TCGA cohort into quartiles based on differential cDC1 signature expression to test dependence of CD8 T:trNK on cDC1 levels (Figure 6G). PDAC patient tumors with the lowest evidence for cDC1 involvement had a weak correlation of trNK_C1 with CD8A and CD8B (p = 0.047; p = 0.045 respectively) which rises to a strong highly significant correlation when the highest levels of cDC1s are present (R = 0.55, trNK_C1 vs CD8A p = 0.0001; R = 0.58, trNK_C1 vs CD8B p = 0.000047) (Figure 6G).

### NK cell signature correlates with improved survival

As the presence of trNK and correlation with CD8 T cells appears boosted by IR, we next further explored our original finding that CD56 correlates with highly significant survival in PDAC (Figure 1, Figure supplement 1). We hypothesized that tumors with a high *NCAM1*/CD56 signature may reflect a greater indication of NK cell recruitment, including a high proportion of trNK (CD56^bright^CD16^low^), whereas tumors with a low CD56 signature might have a lower proportion of infiltrating (tr)NK cells and, therefore, could benefit from IR. Indeed, separating PAAD_TCGA patients into those who received radiotherapy (RTx) versus those that did not, showed that the benefit in overall survival of PDAC patients is only apparent in the CD56^low^ patient group (log rank p <0.0001, Figure 7A). We next explored whether CD56-associated survival is specifically due to the presence of trNK cells using our NK_C1 signature (Table supplement 1). Analysis of primary PDAC tumors from the TCGA (TCGA_PAAD) demonstrates that patients with tumors enriched for the trNK cell (NK_C1) gene signature were associated with improved PDAC survival compared to patients without trNK involvement (Figure 7B, Figure supplement 7A). Similarly to CD56^low^ patients, we find that patients with trNK^low^ benefit from RTx (log rank p <0.005, Figure 7C). Notably, we find that despite an overall poor prognosis, CCL5^high^ patients (TCGA and CPTAC3) are significantly enriched for the trNK_C1 gene signature (Figure 7D), potentially due to strong CCR5 or CCR1-mediated recruitment (Figure supplement 1A). However, as CCL5 also recruits MDSCs, TAMs, Tregs and conventional NK cells via CCR5, CCR5i could prevent a tumor suppressive microenvironment, both directly and indirectly by retaining trNK cells that remove Tregs through FASL (Figure supplement 7B). Therefore, we would expect that patients with CCL5^high^ NK_C1^low^ would perform significantly worse than patients with CCL5^high^NK_C1^high^. Indeed, CCL5^high^ patients enriched for the NK_C1 signature or CD56 had significantly improved overall and disease-free survival (Figure 7E, F). This also supports a model where CCL5 mediated recruitment of NK cells can be beneficial in the absence of CCL5-CCR5 recruitment, likely via CCR1 signaling(Ajuebor et al., 2007) (Figure supplement 7B, 1A). These results suggest that despite high CCL5 levels and an overall poor prognostic outcome, the presence of trNK cells significantly improves survival and provides an opportunity for intervention with the IR+IT combination.

**Figure 7.**
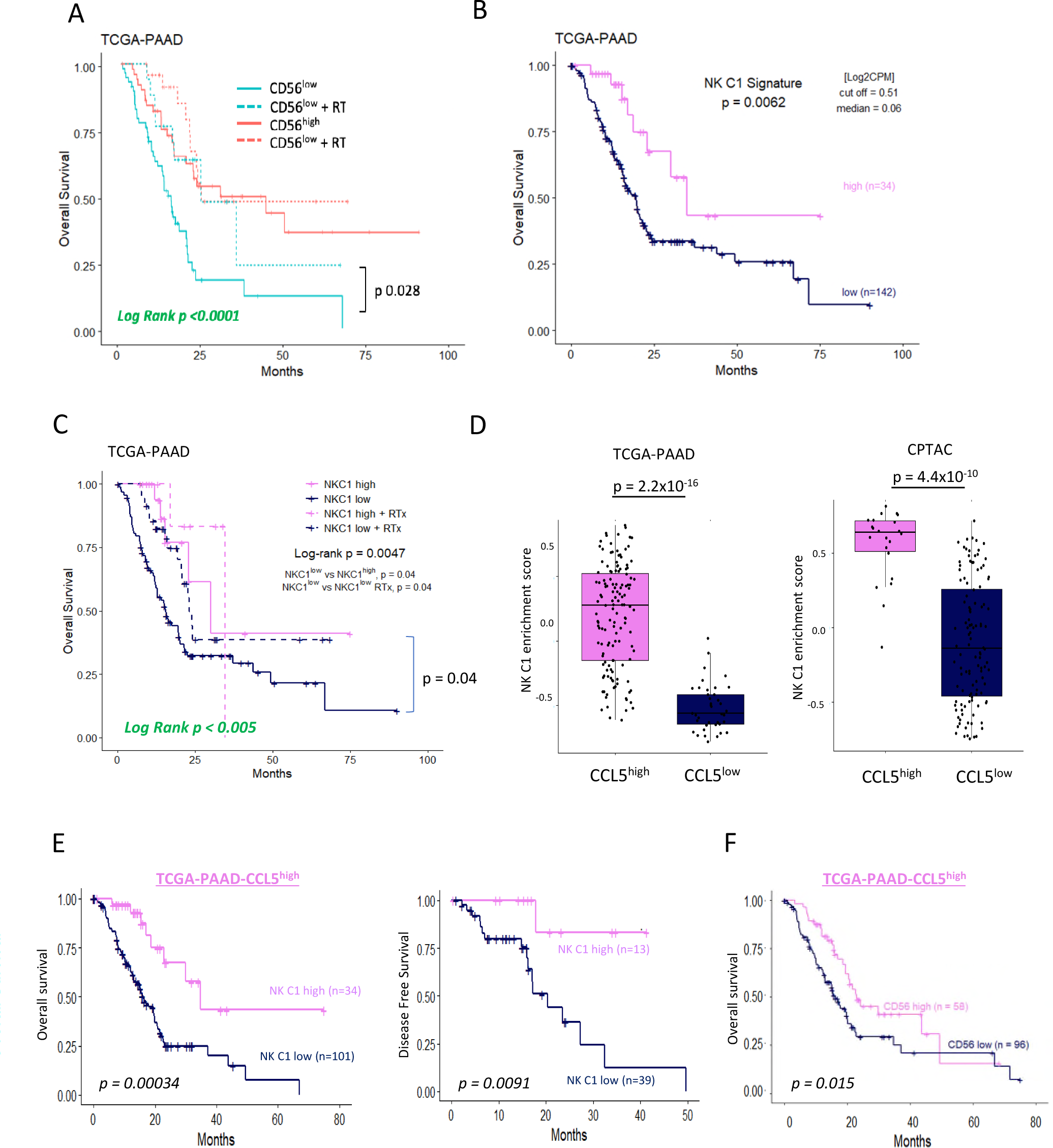
(A) Overall survival analysis correlating CD56 low/high expression with and without radiation therapy in the TCGA dataset. (B) Overall survival analysis correlating the deconvoluted NK C1 signature split into low and high. (C) Overall survival analysis correlating NK C1 low/high enrichment with and without radiation therapy in the TCGA dataset. P-values are adjusted using the Bonferroni method. (D) Boxplot correlating low and high CCL5 expression with NK C1 enrichment score in the TCGA (left) and CPTAC (right) datasets. (E) Overall (left) and disease-free survival (right) analysis of CCL5^high^ patients segregated based on high/low enrichment of trNK (NK_C1) gene signature. (F) Overall survival of CCL5^high^ segregated on CD56 (*NCAM1*) expression.

Finally, we explored whether the NK_C1 gene signature could be a prognostic marker in solid malignancies other than PDAC by expanding our TCGA analysis. The majority of cancers with enriched expression of our NK_C1 gene signature showed improved survival (Figure 8) (apart from endometrium and prostate) as a continuous variable or as a high vs low in a univariate Cox regression (Figure supplement 8A, B). The latter result supports our hypothesis that enrichment of trNK cells is a protective factor across most solid malignancies. Moreover, the ability to enrich for this population with IR+IT combinatorial strategies supports the idea that trNK cells generally improve overall survival though improving CD8 T cell activity in solid cancers.

**Figure 8.**
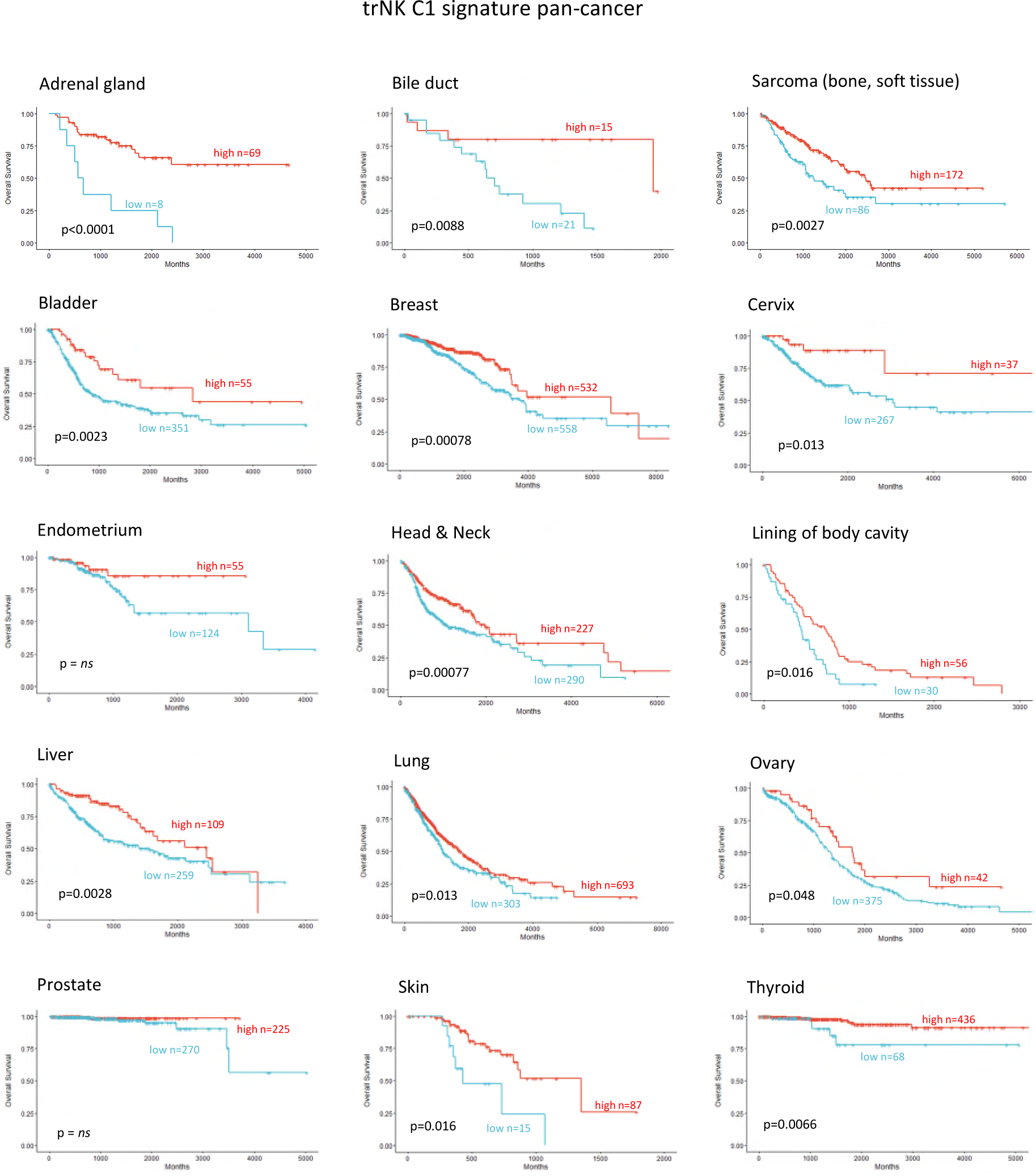
Correlation of trNK (NK_C1) gene signature in human cancer. The trNK cell gene signature is a positive prognostic factor across various malignancies using the TCGA_PAAD dataset. An enrichment score of the NK_C1 gene signature (see Table supplement 1) was first calculated per patient sample in the TCGA RNA-seq dataset using the Gene Set Variation Analysis (GSVA) method. A cut-off value was then determined using the maximally selected rank statistics (max-stat R package) method to divide patients into “high” and “low”.

## Discussion

Novel approaches such as immunotherapies struggle to improve outcome in PDAC as tumors are stromal rich, myeloid-involved immunosuppressive microenvironments devoid of cytotoxic lymphocytes (Chouari et al., 2023). Most strategies to combat immune suppression, e.g. targeting inhibitory checkpoints such as PD1/PDL1, fail as single therapies because PDAC is largely devoid of the CD8 T cells these agents are designed to reactivate. Combining them with inhibitors of myeloid suppression (e.g. CXCR2) to increase CD8 T penetration also failed to impact survival in clinical trials despite showing promise in pre-clinical studies(Steele et al., 2015),(Siolas et al., 2020). Similarly, high-dose hypo-fractionated ablative radiotherapy has been employed to create an acute localized inflammatory response to stimulate intra-tumoral penetration of CD8 T cells, but even in conjunction with PD1/PDL1 blockade this approach has not shown benefit in PDAC (Parikh et al., 2021). Novel strategies in pre-clinical PDAC models comprising ionizing radiation (IR), αPD1 and BMS-687681 (a dual CCR2/CCR5 inhibitor) have shown promising increases in CD8 T cells (Wang et al., 2022) whereas the combination of IR and the bifunctional agent αPD-1/IL-2Rβγ stimulates tumor penetration of polyfunctional stem-like activated CD8 T cells and DNAM1^+^ cytotoxic natural killer (NK) cells(Piper et al., 2023). Together, these approaches indicate potential benefits of a coordinated alteration of the suppressive microenvironment and checkpoint blockade to reduce regulatory T cells (Tregs) and increase NK cell infiltration in addition to supporting CD8 T cell activity. The utility of RT for localized PDAC has been controversial, but innovative technology can now deliver ionizing radiation at higher doses with greater precision(Mills et al., 2022). Rather than simple ablative radiotherapy, this strategy is being employed to increase localized damage that stimulates an acute damage response in immune-cold tumors or increase tumor neoantigens to stimulate adaptive responses (Sodergren et al., 2020). In support of this, the use of IR alongside αPD-1/IL-2Rβγ or FAKi appeared to induce durable immunity in preclinical models (Lander et al., 2022; Piper et al., 2023), suggesting the induction of immunologic memory against tumor antigens is possible.

We were led by our previous clinical trial results implicating serum CCL5 levels as a negative prognostic marker for PDAC survival, which we now validate in two independent validation datasets. CCL5 has both pro-inflammatory roles as a chemoattractant for leukocytes and anti-inflammatory activity via recruitment of CCR5^+^ Tregs (Tan et al., 2009). To maintain beneficial signals but limit pro-tumorigenic signaling, we targeted CCR5 alone using maraviroc. Not surprisingly, in an orthotopic KPC model of PDAC we found that Tregs were restricted by CCR5i directly, and the combination of IR and immunotherapy (IT, CCR5i, αPD1 or CCR5i/αPD1) correlates with a progressive enrichment of CD8 T cells. Intriguingly, CD8 T cells alone were insufficient to explain the loss of cellularity or the increase in necrotic tumors seen with the IR/CCR5i/αPD1 combination, but we do observe a significant increase in total NK and NKT cells that correlates with better local tumor control.

NK infiltration has been associated with positive outcomes in many solid tumors, considered to be due to the positive impact of cytotoxic NK cells in cytotoxic CD8 T-mediated tumor clearance (Nersesian et al., 2021). However, the data supporting the independent contribution of NK activity is difficult to discern due to overlap of expression profiles of cytotoxic cells. Surprisingly, markers of cytotoxic cells or specific CD8/CD4 lymphocyte receptors do not perform as indicators of survival from bulk mRNA-seq datasets (Figure 5). This may be due to sensitivity issues but, more likely, the presence of cytotoxic cells alone does not necessarily indicate beneficial responses in patients (Nersesian et al., 2021). This is supported by the limited efficacy of immunotherapy strategies to tumors with a high mutational burden. NK cells represent a variety of subsets defined by surface markers, found in the periphery (circulating), secondary lymphoid organs (spleen, lymph nodes), and specific tissues (e.g. lung, liver, uterus) where markers of tissue residency increase retention or prevent egress (Hashemi & Malarkannan, 2020). In addition to trNK cells, tumors appear to accumulate NK populations that become less cytotoxic through downregulation of activating receptors, (NKG2D, NKp46, NKp44) and increased expression of inhibitory receptors (NKG2A) or repression/exhaustion markers (e.g. TIGIT, TIM3) (Hashemi & Malarkannan, 2020) (Marcon et al., 2020). Importantly, tissue-resident or hypoactive NK cell subsets commonly display a reduction in cytotoxicity but become highly active immunomodulatory players via expression of XCL1 and XCL2 (de Andrade et al., 2019). In melanoma, NK-mediated expression of XCL1 is crucial for the migration of XCR1^+^ conventional type 1 dendritic cells (cDC1) and therefore the non-cytotoxic NK subsets in tumors may be vital for cDC1-mediated cross-presentation of tumor antigens to CD8 T cells (Böttcher et al., 2018). This may explain why levels of CD8 T cells in tumors or removal of inhibitory signaling is insufficient to gain tumor control. NK-derived FLT3L correlated with increased cDC1 infiltration and improved overall survival and enhanced αPD1 treatment responses in melanoma (Barry et al., 2018), while intratumoral NK cells correlates with increased T-cell and dendritic cell infiltration and improved survival in neuroblastoma (Melaiu et al., 2020). Although our NK_C1 cells are not enriched for *FLT3LG* (data not shown), the data collectively demonstrate that therapeutic strategies aimed at increasing the presence of cDC1 or NK cells may work in combination to support treatments that induce CD8 T cell activity.

Our data suggest that combination therapy to stimulate an acute inflammatory response (IR) together with CCR5i/αPD1 (IT) sufficiently modulates the tumor immune microenvironment to improve tumor control. As frequently observed in solid tumors, reduction of cancer cells correlates with intra-tumoral infiltration of NK cells (CD56^bright^CD16^-^) and improved CD8 T cell penetration(Wu et al., 2020). Surprisingly, our tumors were penetrated by tissue-resident, hypoactive NKG2D^-^ NK cells, suggesting a population with reduced cytotoxicity (Figure 4F, Figure supplement 4) which we correlated to a similar population (NK_C1) identified from scRNA-seq of human PDAC samples (Figure supplement 5C). This population was CD56^+^CD16^-^, in keeping with the apparent beneficial association of CD56 expression with PDAC survival we identified above. NKG2C is a marker of adaptive NK cells during viral infections where they offer a potential memory capability to innate immunity (Brownlie, Scharenberg, Mold, Hard, et al., 2021; Lopez-Botet et al., 2023). Adaptive NK cells were first identified as a subpopulation of blood derived CD16^+^ NK cells in response to viral infection but more recently found to be independent of CD16 expression in lung tissue (Brownlie, Scharenberg, Mold, Hard, et al., 2021). Although we did not specifically examine markers for adaptive NK cells in our mouse model, the expression of NKG2C in the NK_C1 human cluster led us to believe that at least some of these NK cells might be adaptive-like, tissue-resident NK cells similar to those identified in the lung (Brownlie, Scharenberg, Mold, Hård, et al., 2021). Notably, these cells were marked with receptors for tissue residency CD103 (*ITGAE*), CD49a (*ITGA1*), chemokine expression (XCL1/2) and lymphocyte exhaustion markers TIM3 (*HAVCR2*), TIGIT, *TNFRSF4*), potentially suggesting conversion of cytotoxic NK cells to an immunomodulatory phenotype that can persist in tissues (Brownlie, Scharenberg, Mold, Hard, et al., 2021; Ruckert et al., 2022). Single cell RNA-seq from PDAC tissue confirm that trNKs could mediate recruitment of cDC1s via XCL1-XCR1, but communicate additional signals, including IL-16, LIGHT (*TNFSF14*) and MIF-CD74, which have the potential to upregulate MHC-I expression, antigen presentation and co-stimulatory molecules to contribute to cross presentation(Basha et al., 2012; Cruikshank et al., 1987; Zou & Hu, 2005). We also find that trNK may also recruit CD4^+^ Tregs via IL-16 but concomitant FAS-signaling would lead to Treg apoptosis and support tumor control (Figure 6D, Figure supplement 7B).

Strikingly, using our signature we find that elevated involvement of trNKs in PDAC correlates with CD8 T cell recruitment in a cDC1-dependent manner. Moreover, this supports a model where the inactivation of cytotoxic NK cells in tumors appears to be a conversion to immunomodulatory trNKs and perhaps an important mechanistic switch from innate to adaptive immunity. Our therapeutic strategy of short ablative radiation-induced damage (IR) followed by CCR5i/αPD1 (IT) offers a potential regimen to increase disease free survival in PDAC. Likewise, trNK-like cells with similar traits are increasingly being observed across a variety of cancers (Brownlie et al., 2023; Kirchhammer et al., 2022; Marquardt et al., 2015). Finally, our signature identifies patients with a significant survival benefit across 14 tumor types, indicating a universal phenomenon attributable to this hypoactive, tissue-resident, immunomodulatory NK subset.

## Author contributions

Conceptualization, S.M., EO’N. Methodology, S.L., S.H., C.D. Investigation; S.L., S.H., S.G., C.D. E.AJ. Formal analysis, S.G., S.H., C.D., E.AJ. Data curation, S.G., S.L., S.H., C.D., H.F., S.S. Funding acquisition: M.R.M., J.M, E.O’N. Project administration: E.O’N. Supervision: K.J., E.O’N. Writing – original draft, S.G, C.D, E.O’N. Writing – review & editing, S.G., R.BR., G.V., J.M., K.J., E.O’N. All authors read and approved the manuscript.

## Acknowledgements

The authors thank the oncology preclinical imaging core (S.Smart, V.Kersemans, PD.Allen, M. Hill and JM Thompson) for providing contracted small animal MRI and radiotherapy services, Biomedical Services (K.Watson) and H. Xu for technical support. Funding: Kidani Memorial Trust, Pancreatic Cancer UK, Cancer Research UK, Precision Panc (A25233), CRUK Beatson Institute (A31287, A29996) and CRUK Scotland Centre (CTRQQR-2021\100006). This work was supported by Cancer Research UK (CR-UK) grant number CTRQQR-2021\100002, through the Cancer Research UK Oxford Centre.

## References

Ajuebor, M. N., Wondimu, Z., Hogaboam, C. M., Le, T., Proudfoot, A. E., & Swain, M. G. (2007). CCR5 deficiency drives enhanced natural killer cell trafficking to and activation within the liver in murine T cell-mediated hepatitis. Am J Pathol, 170(6), 1975–1988. 10.2353/ajpath.2007.060690

Aldinucci, D., Borghese, C., & Casagrande, N. (2020). The CCL5/CCR5 Axis in Cancer Progression. Cancers (Basel), 12(7). 10.3390/cancers12071765

Alshetaiwi, H., Pervolarakis, N., McIntyre, L. L., Ma, D., Nguyen, Q., Rath, J. A., Nee, K., Hernandez, G., Evans, K., Torosian, L., Silva, A., Walsh, C., & Kessenbrock, K. (2020). Defining the emergence of myeloid-derived suppressor cells in breast cancer using single-cell transcriptomics. Sci Immunol, 5(44). 10.1126/sciimmunol.aay6017

Barry, K. C., Hsu, J., Broz, M. L., Cueto, F. J., Binnewies, M., Combes, A. J., Nelson, A. E., Loo, K., Kumar, R., Rosenblum, M. D., Alvarado, M. D., Wolf, D. M., Bogunovic, D., Bhardwaj, N., Daud, A. I., Ha, P. K., Ryan, W. R., Pollack, J. L., Samad, B., Krummel, M. F. (2018). A natural killer-dendritic cell axis defines checkpoint therapy-responsive tumor microenvironments. Nat Med, 24(8), 1178–1191. 10.1038/s41591-018-0085-8

Basha, G., Omilusik, K., Chavez-Steenbock, A., Reinicke, A. T., Lack, N., Choi, K. B., & Jefferies, W. A. (2012). A CD74-dependent MHC class I endolysosomal cross-presentation pathway. Nat Immunol, 13(3), 237–245. 10.1038/ni.2225

Benkhaled, S., Peters, C., Jullian, N., Arsenijevic, T., Navez, J., Van Gestel, D., Moretti, L., Van Laethem, J. L., & Bouchart, C. (2023). Combination, Modulation and Interplay of Modern Radiotherapy with the Tumor Microenvironment and Targeted Therapies in Pancreatic Cancer: Which Candidates to Boost Radiotherapy? Cancers (Basel), 15(3). 10.3390/cancers15030768

Bialecki, E., Macho Fernandez, E., Ivanov, S., Paget, C., Fontaine, J., Rodriguez, F., Lebeau, L., Ehret, C., Frisch, B., Trottein, F., & Faveeuw, C. (2011). Spleen-resident CD4+ and CD4-CD8alpha-dendritic cell subsets differ in their ability to prime invariant natural killer T lymphocytes. PLoS One, 6(10), e26919. 10.1371/journal.pone.0026919

Bockorny, B., Semenisty, V., Macarulla, T., Borazanci, E., Wolpin, B. M., Stemmer, S. M., Golan, T., Geva, R., Borad, M. J., Pedersen, K. S., Park, J. O., Ramirez, R. A., Abad, D. G., Feliu, J., Munoz, A., Ponz-Sarvise, M., Peled, A., Lustig, T. M., Bohana-Kashtan, O., Hidalgo, M. (2020). BL-8040, a CXCR4 antagonist, in combination with pembrolizumab and chemotherapy for pancreatic cancer: the COMBAT trial. Nat Med, 26(6), 878–885. 10.1038/s41591-020-0880-x

Böttcher, J. P., Bonavita, E., Chakravarty, P., Blees, H., Cabeza-Cabrerizo, M., Sammicheli, S., Rogers, N. C., Sahai, E., Zelenay, S., & Reis e Sousa, C. (2018). NK Cells Stimulate Recruitment of cDC1 into the Tumor Microenvironment Promoting Cancer Immune Control. Cell, 172(5), 1022–1037.e1014. 10.1016/j.cell.2018.01.004

Brownlie, D., Scharenberg, M., Mold, J. E., Hard, J., Kekalainen, E., Buggert, M., Nguyen, S., Wilson, J. N., Al-Ameri, M., Ljunggren, H. G., Marquardt, N., & Michaelsson, J. (2021). Expansions of adaptive-like NK cells with a tissue-resident phenotype in human lung and blood. Proc Natl Acad Sci U S A, 118(11). 10.1073/pnas.2016580118

Brownlie, D., Scharenberg, M., Mold, J. E., Hård, J., Kekäläinen, E., Buggert, M., Nguyen, S., Wilson, J. N., Al-Ameri, M., Ljunggren, H. G., Marquardt, N., & Michaëlsson, J. (2021). Expansions of adaptive-like NK cells with a tissue-resident phenotype in human lung and blood. Proc Natl Acad Sci U S A, 118(11). 10.1073/pnas.2016580118

Brownlie, D., von Kries, A., Valenzano, G., Wild, N., Yilmaz, E., Safholm, J., Al-Ameri, M., Alici, E., Ljunggren, H. G., Schliemann, I., Aricak, O., Haglund de Flon, F., Michaelsson, J., & Marquardt, N. (2023). Accumulation of tissue-resident natural killer cells, innate lymphoid cells, and CD8(+) T cells towards the center of human lung tumors. Oncoimmunology, 12(1), 2233402. 10.1080/2162402X.2023.2233402

Bryceson, Y. T., & Ljunggren, H. G. (2008). Tumor cell recognition by the NK cell activating receptor NKG2D. Eur J Immunol, 38(11), 2957–2961. 10.1002/eji.200838833

Cao, L., Huang, C., Cui Zhou, D., Hu, Y., Lih, T. M., Savage, S. R., Krug, K., Clark, D. J., Schnaubelt, M., Chen, L., da Veiga Leprevost, F., Eguez, R. V., Yang, W., Pan, J., Wen, B., Dou, Y., Jiang, W., Liao, Y., Shi, Z., Clinical Proteomic Tumor Analysis, C. (2021). Proteogenomic characterization of pancreatic ductal adenocarcinoma. Cell, 184(19), 5031–5052 e5026. 10.1016/j.cell.2021.08.023

Caushi, J. X., Zhang, J., Ji, Z., Vaghasia, A., Zhang, B., Hsiue, E. H., Mog, B. J., Hou, W., Justesen, S., Blosser, R., Tam, A., Anagnostou, V., Cottrell, T. R., Guo, H., Chan, H. Y., Singh, D., Thapa, S., Dykema, A. G., Burman, P., Smith, K. N. (2021). Transcriptional programs of neoantigen-specific TIL in anti-PD-1-treated lung cancers. Nature, 596(7870), 126–132. 10.1038/s41586-021-03752-4

Chen, I. M., Johansen, J. S., Theile, S., Hjaltelin, J. X., Novitski, S. I., Brunak, S., Hasselby, J. P., Willemoe, G. L., Lorentzen, T., Madsen, K., Jensen, B. V., Wilken, E. E., Geertsen, P., Behrens, C., Nolsoe, C., Hermann, K. L., Svane, I. M., & Nielsen, D. (2022). Randomized Phase II Study of Nivolumab With or Without Ipilimumab Combined With Stereotactic Body Radiotherapy for Refractory Metastatic Pancreatic Cancer (CheckPAC). J Clin Oncol, 40(27), 3180–3189. 10.1200/jco.21.02511

Chouari, T., La Costa, F. S., Merali, N., Jessel, M. D., Sivakumar, S., Annels, N., & Frampton, A. E. (2023). Advances in Immunotherapeutics in Pancreatic Ductal Adenocarcinoma. Cancers (Basel), 15(17). 10.3390/cancers15174265

Cruikshank, W. W., Berman, J. S., Theodore, A. C., Bernardo, J., & Center, D. M. (1987). Lymphokine activation of T4+ T lymphocytes and monocytes. J Immunol, 138(11), 3817–3823. https://www.ncbi.nlm.nih.gov/pubmed/3108375

de Andrade, L. F., Lu, Y., Luoma, A., Ito, Y., Pan, D., Pyrdol, J. W., Yoon, C. H., Yuan, G. C., & Wucherpfennig, K. W. (2019). Discovery of specialized NK cell populations infiltrating human melanoma metastases. JCI Insight, 4(23). 10.1172/jci.insight.133103

Di Pilato, M., Kfuri-Rubens, R., Pruessmann, J. N., Ozga, A. J., Messemaker, M., Cadilha, B. L., Sivakumar, R., Cianciaruso, C., Warner, R. D., Marangoni, F., Carrizosa, E., Lesch, S., Billingsley, J., Perez-Ramos, D., Zavala, F., Rheinbay, E., Luster, A. D., Gerner, M. Y., Kobold, S.., Mempel, T. R. (2021). CXCR6 positions cytotoxic T cells to receive critical survival signals in the tumor microenvironment. Cell, 184(17), 4512–4530 e4522. 10.1016/j.cell.2021.07.015

Doi, T., Muro, K., Ishii, H., Kato, T., Tsushima, T., Takenoyama, M., Oizumi, S., Gemmoto, K., Suna, H., Enokitani, K., Kawakami, T., Nishikawa, H., & Yamamoto, N. (2019). A Phase I Study of the Anti-CC Chemokine Receptor 4 Antibody, Mogamulizumab, in Combination with Nivolumab in Patients with Advanced or Metastatic Solid Tumors. Clin Cancer Res, 25(22), 6614–6622. 10.1158/1078-0432.CCR-19-1090

Groh, V., Wu, J., Yee, C., & Spies, T. (2002). Tumour-derived soluble MIC ligands impair expression of NKG2D and T-cell activation. Nature, 419(6908), 734–738. 10.1038/nature01112

Hashemi, E., & Malarkannan, S. (2020). Tissue-Resident NK Cells: Development, Maturation, and Clinical Relevance. Cancers (Basel), 12(6). 10.3390/cancers12061553

Hemmatazad, H., & Berger, M. D. (2021). CCR5 is a potential therapeutic target for cancer. Expert Opin Ther Targets, 25(4), 311–327. 10.1080/14728222.2021.1902505

Huang, H., Zepp, M., Georges, R. B., Jarahian, M., Kazemi, M., Eyol, E., & Berger, M. R. (2020). The CCR5 antagonist maraviroc causes remission of pancreatic cancer liver metastasis in nude rats based on cell cycle inhibition and apoptosis induction. Cancer Lett, 474, 82–93. 10.1016/j.canlet.2020.01.009

Jin, S., Guerrero-Juarez, C. F., Zhang, L., Chang, I., Ramos, R., Kuan, C. H., Myung, P., Plikus, M. V., & Nie, Q. (2021). Inference and analysis of cell-cell communication using CellChat. Nat Commun, 12(1), 1088. 10.1038/s41467-021-21246-9

Kaiser, B. K., Yim, D., Chow, I. T., Gonzalez, S., Dai, Z., Mann, H. H., Strong, R. K., Groh, V., & Spies, T. (2007). Disulphide-isomerase-enabled shedding of tumour-associated NKG2D ligands. Nature, 447(7143), 482–486. 10.1038/nature05768

Kaser, A., Dunzendorfer, S., Offner, F. A., Ryan, T., Schwabegger, A., Cruikshank, W. W., Wiedermann, C. J., & Tilg, H. (1999). A role for IL-16 in the cross-talk between dendritic cells and T cells. J Immunol, 163(6), 3232–3238.

Kirchhammer, N., Trefny, M. P., Natoli, M., Brucher, D., Smith, S. N., Werner, F., Koch, V., Schreiner, D., Bartoszek, E., Buchi, M., Schmid, M., Breu, D., Hartmann, K. P., Zaytseva, P., Thommen, D. S., Laubli, H., Bottcher, J. P., Stanczak, M. A., Kashyap, A. S., Zippelius, A. (2022). NK cells with tissue-resident traits shape response to immunotherapy by inducing adaptive antitumor immunity. Sci Transl Med, 14(653), eabm9043. 10.1126/scitranslmed.abm9043

Lander, V. E., Belle, J. I., Kingston, N. L., Herndon, J. M., Hogg, G. D., Liu, X., Kang, L. I., Knolhoff, B. L., Bogner, S. J., Baer, J. M., Zuo, C., Borcherding, N. C., Lander, D. P., Mpoy, C., Scott, J., Zahner, M., Rogers, B. E., Schwarz, J. K., Kim, H., & DeNardo, D. G. (2022). Stromal Reprogramming by FAK Inhibition Overcomes Radiation Resistance to Allow for Immune Priming and Response to Checkpoint Blockade. Cancer Discov, 12(12), 2774–2799. 10.1158/2159-8290.CD-22-0192

Lopez-Botet, M., De Maria, A., Muntasell, A., Della Chiesa, M., & Vilches, C. (2023). Adaptive NK cell response to human cytomegalovirus: Facts and open issues. Semin Immunol, 65, 101706. 10.1016/j.smim.2022.101706

Mantovani, A., Marchesi, F., Malesci, A., Laghi, L., & Allavena, P. (2017). Tumour-associated macrophages as treatment targets in oncology. Nat Rev Clin Oncol, 14(7), 399–416. 10.1038/nrclinonc.2016.217

Marcon, F., Zuo, J., Pearce, H., Nicol, S., Margielewska-Davies, S., Farhat, M., Mahon, B., Middleton, G., Brown, R., Roberts, K. J., & Moss, P. (2020). NK cells in pancreatic cancer demonstrate impaired cytotoxicity and a regulatory IL-10 phenotype. Oncoimmunology, 9(1), 1845424. 10.1080/2162402X.2020.1845424

Marquardt, N., Beziat, V., Nystrom, S., Hengst, J., Ivarsson, M. A., Kekalainen, E., Johansson, H., Mjosberg, J., Westgren, M., Lankisch, T. O., Wedemeyer, H., Ellis, E. C., Ljunggren, H. G., Michaelsson, J., & Bjorkstrom, N. K. (2015). Cutting edge: identification and characterization of human intrahepatic CD49a+ NK cells. J Immunol, 194(6), 2467–2471. 10.4049/jimmunol.1402756

Matzke-Ogi, A., Jannasch, K., Shatirishvili, M., Fuchs, B., Chiblak, S., Morton, J., Tawk, B., Lindner, T., Sansom, O., Alves, F., Warth, A., Schwager, C., Mier, W., Kleeff, J., Ponta, H., Abdollahi, A., & Orian-Rousseau, V. (2016). Inhibition of Tumor Growth and Metastasis in Pancreatic Cancer Models by Interference With CD44v6 Signaling. Gastroenterology, 150(2), 513–525 e510. 10.1053/j.gastro.2015.10.020

McFadden, C., Morgan, R., Rahangdale, S., Green, D., Yamasaki, H., Center, D., & Cruikshank, W. (2007). Preferential migration of T regulatory cells induced by IL-16. J Immunol, 179(10), 6439–6445. 10.4049/jimmunol.179.10.6439

Melaiu, O., Chierici, M., Lucarini, V., Jurman, G., Conti, L. A., De Vito, R., Boldrini, R., Cifaldi, L., Castellano, A., Furlanello, C., Barnaba, V., Locatelli, F., & Fruci, D. (2020). Cellular and gene signatures of tumor-infiltrating dendritic cells and natural-killer cells predict prognosis of neuroblastoma. Nat Commun, 11(1), 5992. 10.1038/s41467-020-19781-y

Mijnheer, G., Lutter, L., Mokry, M., van der Wal, M., Scholman, R., Fleskens, V., Pandit, A., Tao, W., Wekking, M., Vervoort, S., Roberts, C., Petrelli, A., Peeters, J. G. C., Knijff, M., de Roock, S., Vastert, S., Taams, L. S., van Loosdregt, J., & van Wijk, F. (2021). Conserved human effector Treg cell transcriptomic and epigenetic signature in arthritic joint inflammation. Nat Commun, 12(1), 2710. 10.1038/s41467-021-22975-7

Mills, B. N., Qiu, H., Drage, M. G., Chen, C., Mathew, J. S., Garrett-Larsen, J., Ye, J., Uccello, T. P., Murphy, J. D., Belt, B. A., Lord, E. M., Katz, A. W., Linehan, D. C., & Gerber, S. A. (2022). Modulation of the Human Pancreatic Ductal Adenocarcinoma Immune Microenvironment by Stereotactic Body Radiotherapy. Clin Cancer Res, 28(1), 150–162. 10.1158/1078-0432.CCR-21-2495

Mohindra, N. A., Kircher, S. M., Nimeiri, H. S., Benson, A. B., Rademaker, A., Alonso, E., Blatner, N., Khazaie, K., & Mulcahy, M. F. (2015). Results of the phase Ib study of ipilimumab and gemcitabine for advanced pancreas cancer. Journal of Clinical Oncology, 33(15_suppl), e15281–e15281. 10.1200/jco.2015.33.15_suppl.e15281

Monti, P., Marchesi, F., Reni, M., Mercalli, A., Sordi, V., Zerbi, A., Balzano, G., Di Carlo, V., Allavena, P., & Piemonti, L. (2004). A comprehensive in vitro characterization of pancreatic ductal carcinoma cell line biological behavior and its correlation with the structural and genetic profile. Virchows Arch, 445(3), 236–247. 10.1007/s00428-004-1053-x

Nersesian, S., Schwartz, S. L., Grantham, S. R., MacLean, L. K., Lee, S. N., Pugh-Toole, M., & Boudreau, J. E. (2021). NK cell infiltration is associated with improved overall survival in solid cancers: A systematic review and meta-analysis. Transl Oncol, 14(1), 100930. 10.1016/j.tranon.2020.100930

Parikh, A. R., Szabolcs, A., Allen, J. N., Clark, J. W., Wo, J. Y., Raabe, M., Thel, H., Hoyos, D., Mehta, A., Arshad, S., Lieb, D. J., Drapek, L. C., Blaszkowsky, L. S., Giantonio, B. J., Weekes, C. D., Zhu, A. X., Goyal, L., Nipp, R. D., Dubois, J. S., Hong, T. S. (2021). Radiation therapy enhances immunotherapy response in microsatellite stable colorectal and pancreatic adenocarcinoma in a phase II trial. Nat Cancer, 2(11), 1124–1135. 10.1038/s43018-021-00269-7

Peng, Y. P., Zhu, Y., Zhang, J. J., Xu, Z. K., Qian, Z. Y., Dai, C. C., Jiang, K. R., Wu, J. L., Gao, W. T., Li, Q., Du, Q., & Miao, Y. (2013). Comprehensive analysis of the percentage of surface receptors and cytotoxic granules positive natural killer cells in patients with pancreatic cancer, gastric cancer, and colorectal cancer. J Transl Med, 11, 262. 10.1186/1479-5876-11-262

Piper, M., Hoen, M., Darragh, L. B., Knitz, M. W., Nguyen, D., Gadwa, J., Durini, G., Karakoc, I., Grier, A., Neupert, B., Van Court, B., Abdelazeem, K. N. M., Yu, J., Olimpo, N. A., Corbo, S., Ross, R. B., Pham, T. T., Joshi, M., Kedl, R. M., Karam, S. D. (2023). Simultaneous targeting of PD-1 and IL-2Rbetagamma with radiation therapy inhibits pancreatic cancer growth and metastasis. Cancer Cell, 41(5), 950–969 e956. 10.1016/j.ccell.2023.04.001

Poli, A., Michel, T., Theresine, M., Andres, E., Hentges, F., & Zimmer, J. (2009). CD56bright natural killer (NK) cells: an important NK cell subset. Immunology, 126(4), 458–465. 10.1111/j.1365-2567.2008.03027.x

Quatrini, L., Molfetta, R., Zitti, B., Peruzzi, G., Fionda, C., Capuano, C., Galandrini, R., Cippitelli, M., Santoni, A., & Paolini, R. (2015). Ubiquitin-dependent endocytosis of NKG2D-DAP10 receptor complexes activates signaling and functions in human NK cells. Sci Signal, 8(400), ra108. 10.1126/scisignal.aab2724

Rand, T. H., Cruikshank, W. W., Center, D. M., & Weller, P. F. (1991). CD4-mediated stimulation of human eosinophils: lymphocyte chemoattractant factor and other CD4-binding ligands elicit eosinophil migration. J Exp Med, 173(6), 1521–1528. 10.1084/jem.173.6.1521

Royal, R. E., Levy, C., Turner, K., Mathur, A., Hughes, M., Kammula, U. S., Sherry, R. M., Topalian, S. L., Yang, J. C., Lowy, I., & Rosenberg, S. A. (2010). Phase 2 trial of single agent Ipilimumab (anti-CTLA-4) for locally advanced or metastatic pancreatic adenocarcinoma. J Immunother, 33(8), 828–833. 10.1097/CJI.0b013e3181eec14c

Ruckert, T., Lareau, C. A., Mashreghi, M. F., Ludwig, L. S., & Romagnani, C. (2022). Clonal expansion and epigenetic inheritance of long-lasting NK cell memory. Nat Immunol, 23(11), 1551–1563. 10.1038/s41590-022-01327-7

Simhadri, V. R., Reiners, K. S., Hansen, H. P., Topolar, D., Simhadri, V. L., Nohroudi, K., Kufer, T. A., Engert, A., & Pogge von Strandmann, E. (2008). Dendritic cells release HLA-B-associated transcript-3 positive exosomes to regulate natural killer function. PLoS One, 3(10), e3377. 10.1371/journal.pone.0003377

Singh, S. K., Mishra, M. K., Eltoum, I. A., Bae, S., Lillard, J. W., Jr., & Singh, R. (2018). CCR5/CCL5 axis interaction promotes migratory and invasiveness of pancreatic cancer cells. Sci Rep, 8(1), 1323. 10.1038/s41598-018-19643-0

Siolas, D., Morrissey, C., & Oberstein, P. E. (2020). The Achilles’ Heel of Pancreatic Cancer: Targeting pancreatic cancer’s unique immunologic characteristics and metabolic dependencies in clinical trials. J Pancreatol, 3(3), 121–131. 10.1097/JP9.0000000000000052

Smith, S. L., Kennedy, P. R., Stacey, K. B., Worboys, J. D., Yarwood, A., Seo, S., Solloa, E. H., Mistretta, B., Chatterjee, S. S., Gunaratne, P., Allette, K., Wang, Y. C., Smith, M. L., Sebra, R., Mace, E. M., Horowitz, A., Thomson, W., Martin, P., Eyre, S., & Davis, D. M. (2020). Diversity of peripheral blood human NK cells identified by single-cell RNA sequencing. Blood Adv, 4(7), 1388–1406. 10.1182/bloodadvances.2019000699

Sodergren, M. H., Mangal, N., Wasan, H., Sadanandam, A., Balachandran, V. P., Jiao, L. R., & Habib, N. (2020). Immunological combination treatment holds the key to improving survival in pancreatic cancer. J Cancer Res Clin Oncol, 146(11), 2897–2911. 10.1007/s00432-020-03332-5

Steele, C. W., Karim, S. A., Foth, M., Rishi, L., Leach, J. D., Porter, R. J., Nixon, C., Jeffry Evans, T. R., Carter, C. R., Nibbs, R. J., Sansom, O. J., & Morton, J. P. (2015). CXCR2 inhibition suppresses acute and chronic pancreatic inflammation. J Pathol, 237(1), 85–97. 10.1002/path.4555

Steele, N. G., Carpenter, E. S., Kemp, S. B., Sirihorachai, V. R., The, S., Delrosario, L., Lazarus, J., Amir, E. D., Gunchick, V., Espinoza, C., Bell, S., Harris, L., Lima, F., Irizarry-Negron, V., Paglia, D., Macchia, J., Chu, A. K. Y., Schofield, H., Wamsteker, E. J., Pasca di Magliano, M. (2020). Multimodal Mapping of the Tumor and Peripheral Blood Immune Landscape in Human Pancreatic Cancer. Nat Cancer, 1(11), 1097–1112. 10.1038/s43018-020-00121-4

Tan, M. C., Goedegebuure, P. S., Belt, B. A., Flaherty, B., Sankpal, N., Gillanders, W. E., Eberlein, T. J., Hsieh, C. S., & Linehan, D. C. (2009). Disruption of CCR5-dependent homing of regulatory T cells inhibits tumor growth in a murine model of pancreatic cancer. J Immunol, 182(3), 1746–1755. 10.4049/jimmunol.182.3.1746

Uhlen, M., Zhang, C., Lee, S., Sjostedt, E., Fagerberg, L., Bidkhori, G., Benfeitas, R., Arif, M., Liu, Z., Edfors, F., Sanli, K., von Feilitzen, K., Oksvold, P., Lundberg, E., Hober, S., Nilsson, P., Mattsson, J., Schwenk, J. M., Brunnstrom, H., Ponten, F. (2017). A pathology atlas of the human cancer transcriptome. Science, 357(6352). 10.1126/science.aan2507

Vella, J. L., Molodtsov, A., Angeles, C. V., Branchini, B. R., Turk, M. J., & Huang, Y. H. (2021). Dendritic cells maintain anti-tumor immunity by positioning CD8 skin-resident memory T cells. Life Sci Alliance, 4(10). 10.26508/lsa.202101056

Vremec, D., Pooley, J., Hochrein, H., Wu, L., & Shortman, K. (2000). CD4 and CD8 expression by dendritic cell subtypes in mouse thymus and spleen. J Immunol, 164(6), 2978–2986. 10.4049/jimmunol.164.6.2978

Wang, J., Saung, M. T., Li, K., Fu, J., Fujiwara, K., Niu, N., Muth, S., Wang, J., Xu, Y., Rozich, N., Zlomke, H., Chen, S., Espinoza, B., Henderson, M., Funes, V., Herbst, B., Ding, D., Twyman-Saint Victor, C., Zhao, Q., Zheng, L. (2022). CCR2/CCR5 inhibitor permits the radiation-induced effector T cell infiltration in pancreatic adenocarcinoma. J Exp Med, 219(5). 10.1084/jem.20211631

Wang, L., He, T., Liu, J., Tai, J., Wang, B., Chen, Z., & Quan, Z. (2021). Pan-cancer analysis reveals tumor-associated macrophage communication in the tumor microenvironment. Exp Hematol Oncol, 10(1), 31. 10.1186/s40164-021-00226-1

Weiss, G. J., Blaydorn, L., Beck, J., Bornemann-Kolatzki, K., Urnovitz, H., Schutz, E., & Khemka, V. (2018). Phase Ib/II study of gemcitabine, nab-paclitaxel, and pembrolizumab in metastatic pancreatic adenocarcinoma. Invest New Drugs, 36(1), 96–102. 10.1007/s10637-017-0525-1

Weiss, G. J., Waypa, J., Blaydorn, L., Coats, J., McGahey, K., Sangal, A., Niu, J., Lynch, C. A., Farley, J. H., & Khemka, V. (2017). A phase Ib study of pembrolizumab plus chemotherapy in patients with advanced cancer (PembroPlus). Br J Cancer, 117(1), 33–40. 10.1038/bjc.2017.145

Willenbrock, F., Cox, C. M., Parkes, E. E., Wilhelm-Benartzi, C. S., Abraham, A. G., Owens, R., Sabbagh, A., Jones, C. M., Hughes, D. L. I., Maughan, T., Hurt, C. N., O’Neill, E. E., & Mukherjee, S. (2021). Circulating biomarkers and outcomes from a randomised phase 2 trial of gemcitabine versus capecitabine-based chemoradiotherapy for pancreatic cancer. Br J Cancer, 124(3), 581–586. 10.1038/s41416-020-01120-z

Wu, S. Y., Fu, T., Jiang, Y. Z., & Shao, Z. M. (2020). Natural killer cells in cancer biology and therapy. Mol Cancer, 19(1), 120. 10.1186/s12943-020-01238-x

Zou, G. M., & Hu, W. Y. (2005). LIGHT regulates CD86 expression on dendritic cells through NF-kappaB, but not JNK/AP-1 signal transduction pathway. J Cell Physiol, 205(3), 437–443. 10.1002/jcp.20420

