## Supplementary material for "Tissue-resident NK cells support survival in pancreatic cancer through promotion of cDC1-CD8T activity": Figure Supplement File

Figure supplement 1

A

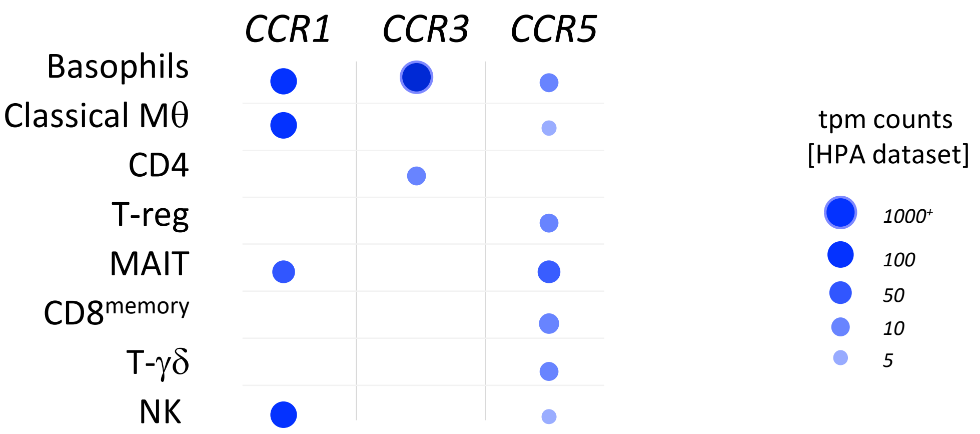


B

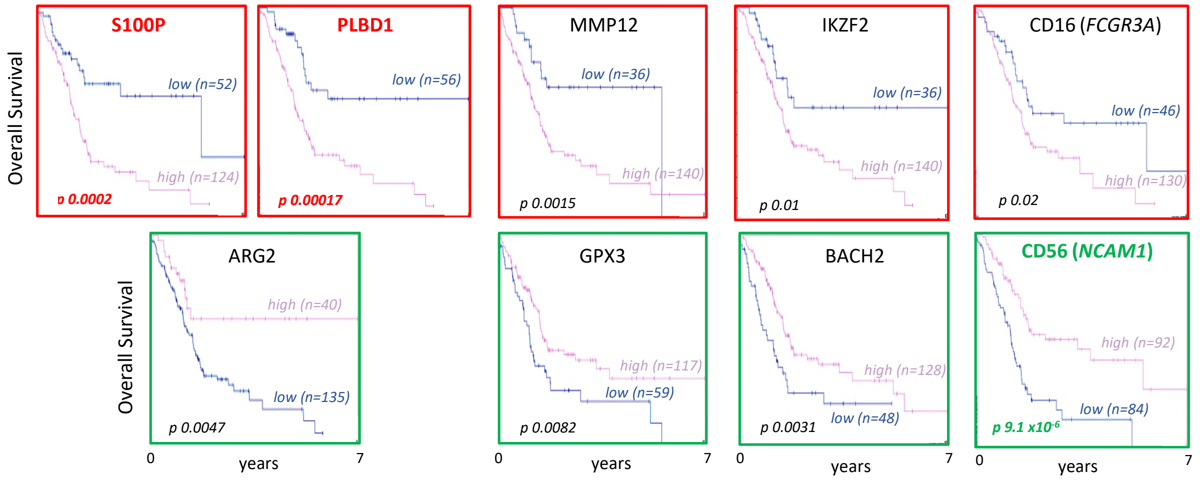


C

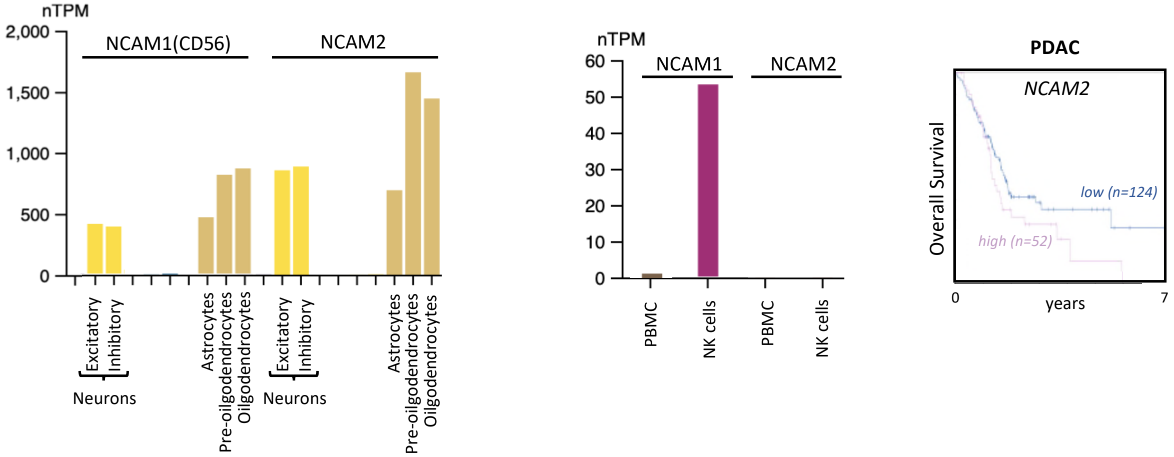

Figure supplement 2

A

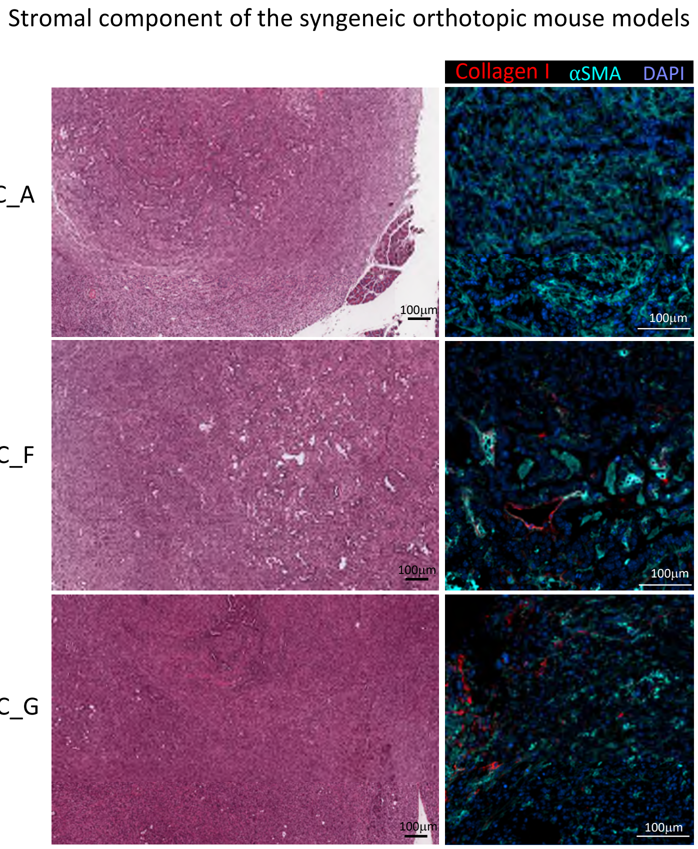


B

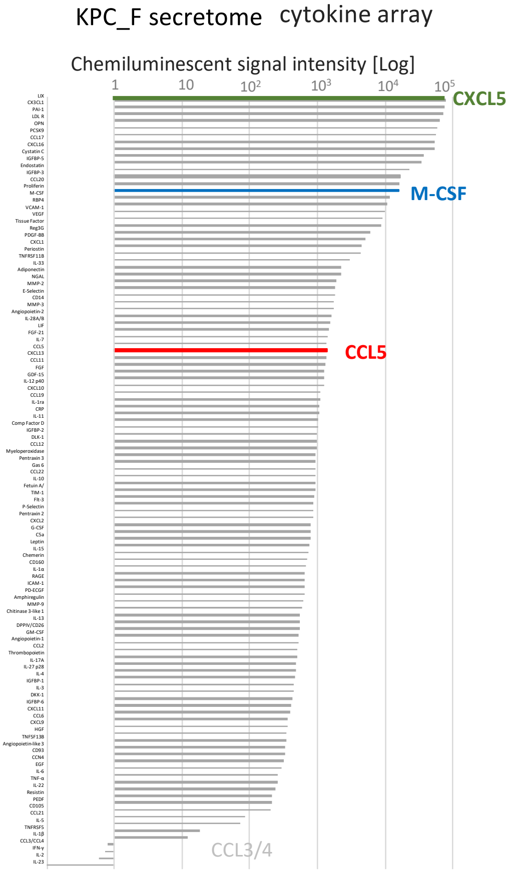


C

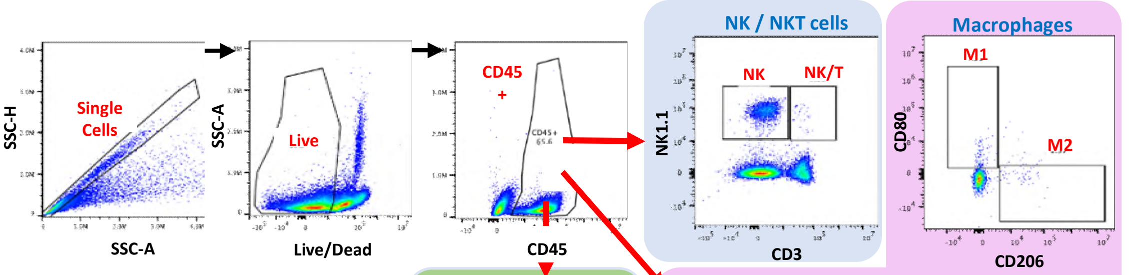


| Antigen | Fluorophore |
|---|---|
| CD45 | AF532 |
| CD19 | SB436 |
| **CD3** | eF450 |
| **NK1.1** | BV650 |
| **CD8a** | BV570 |
| **CD4** | BV510 |
| CD25 | AF647 |
| FOXP3 | PE |
| CD206 | BV785 |
| F4/80 | BV480 |
| CD80 | APC |
| CD279 (PD-1) | PE-Cy7 |
| CD195 (CCR5) | VB-FITC |
| CD11b | AF700 |
| Ly6C | PE-Dazzle |
| Ly6G | BV421 |

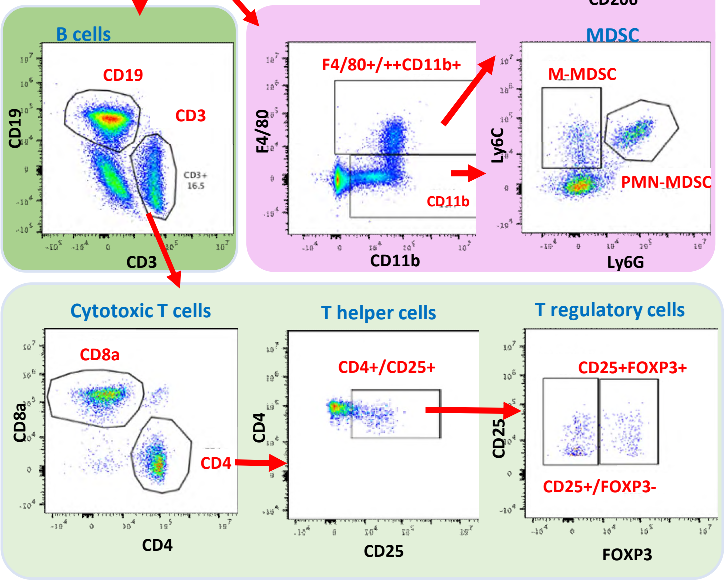

Figure supplement 3

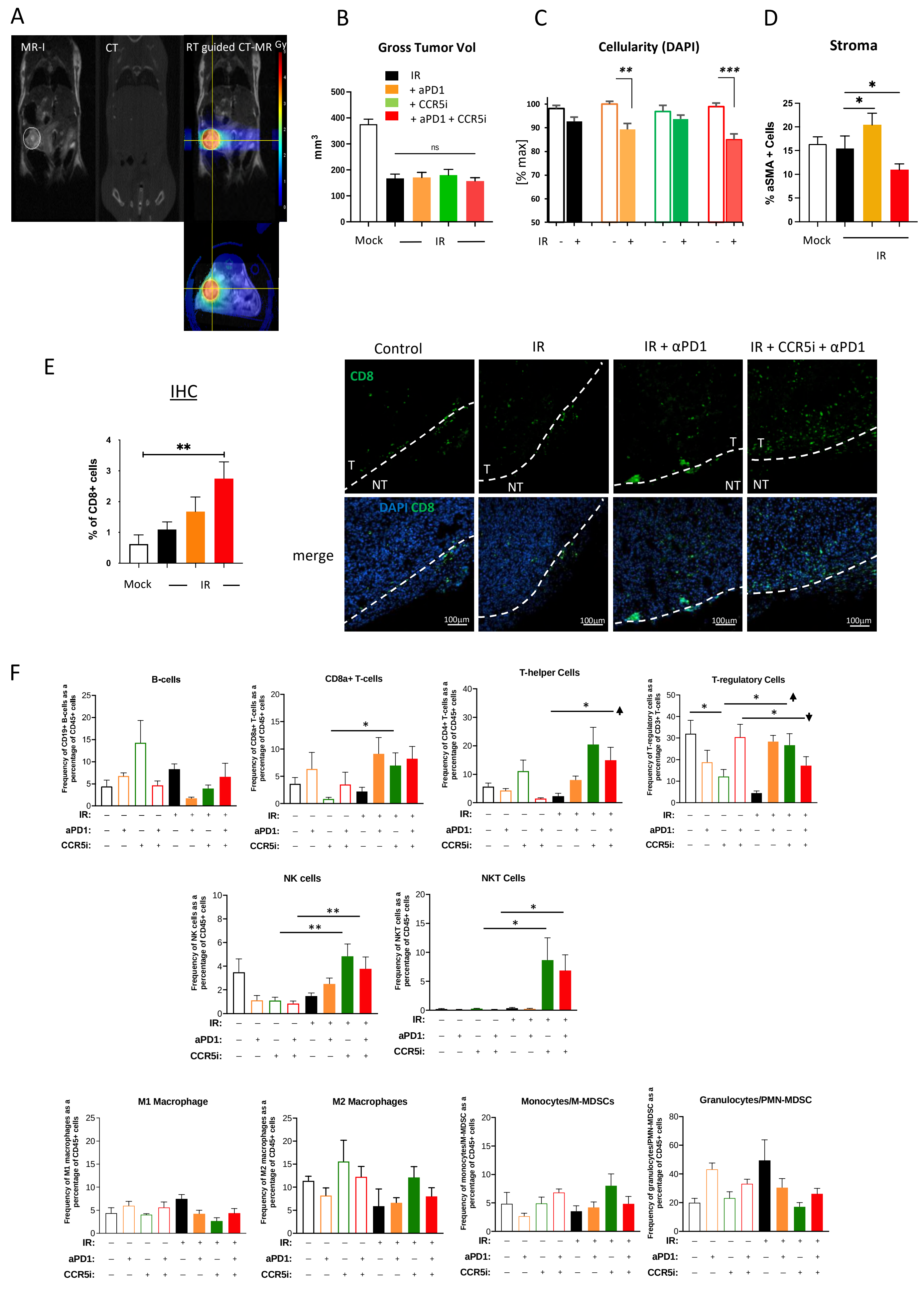

Figure supplement 4

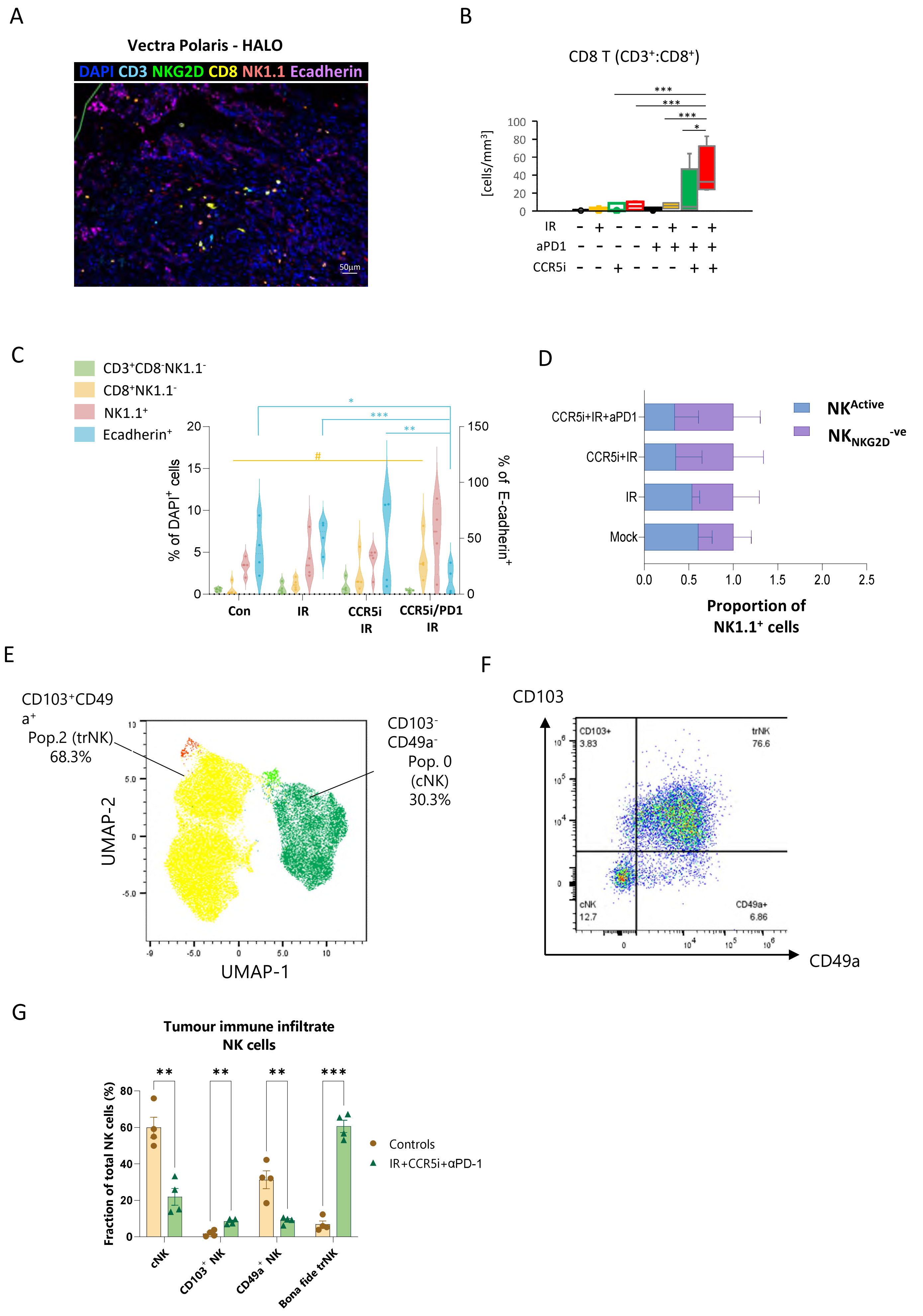

Figure supplement 5

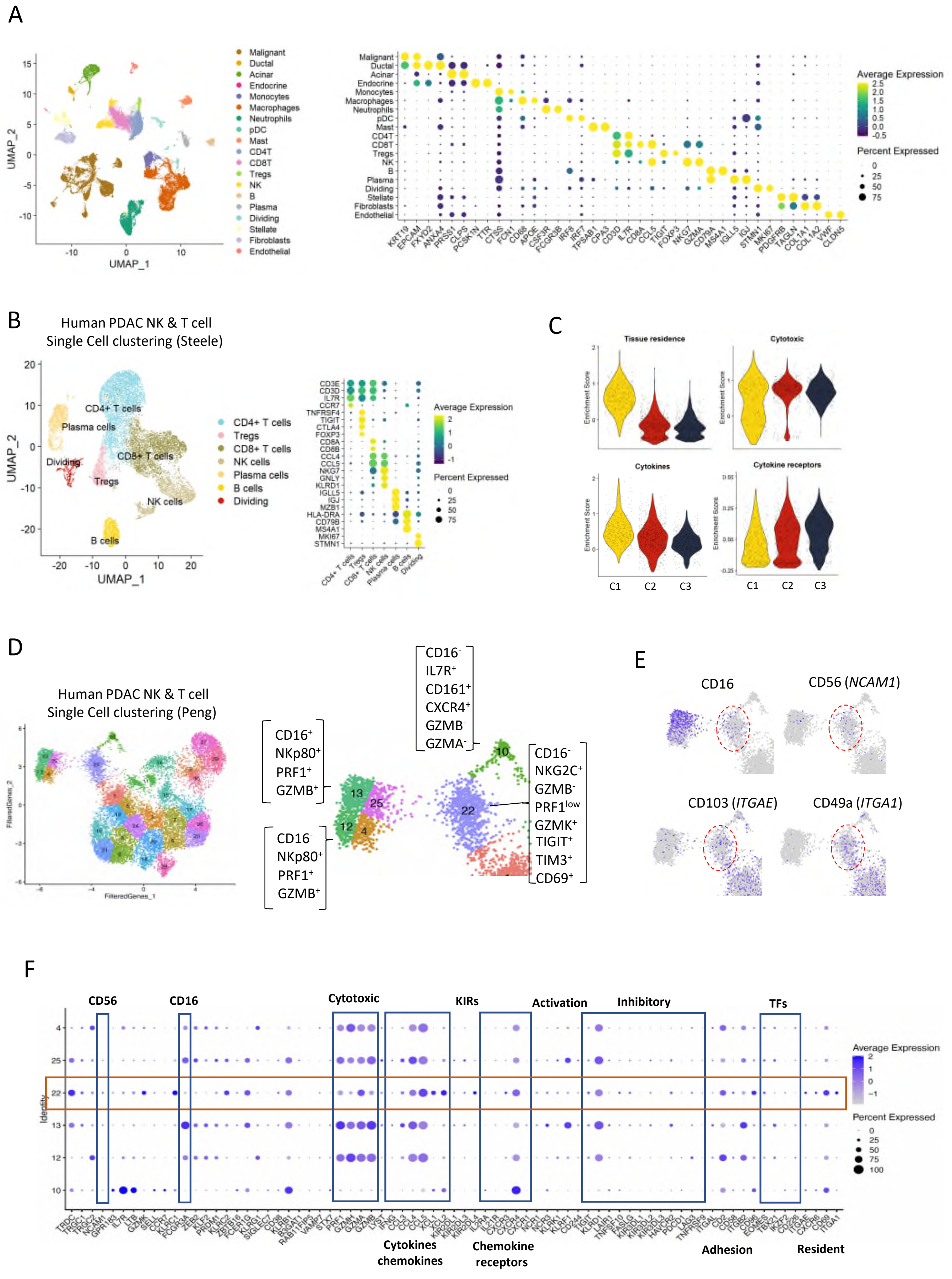

Figure supplement 6

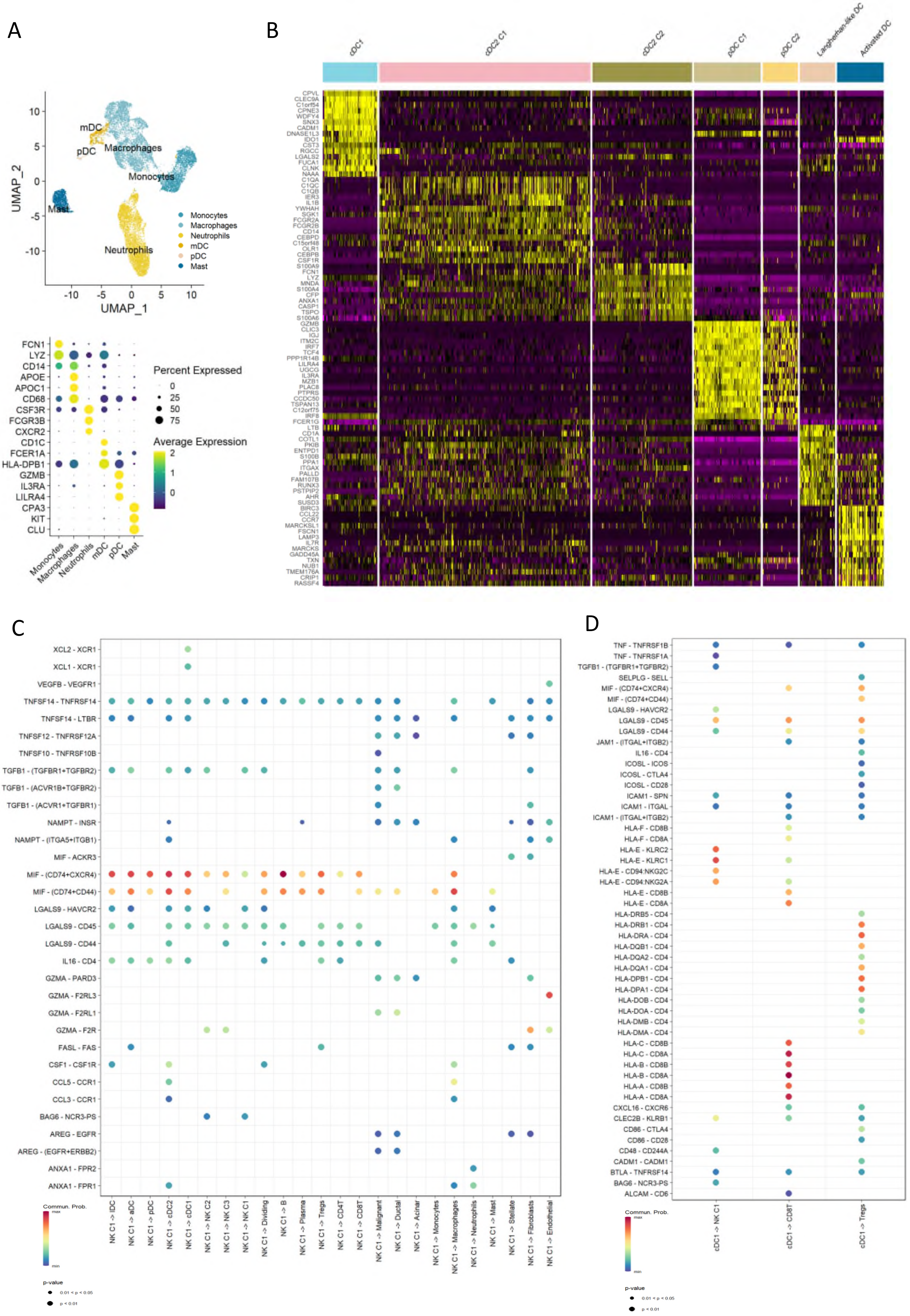

Figure supplement 7

A

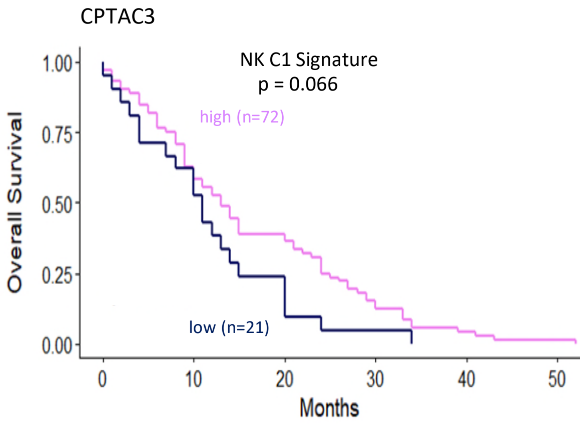


B

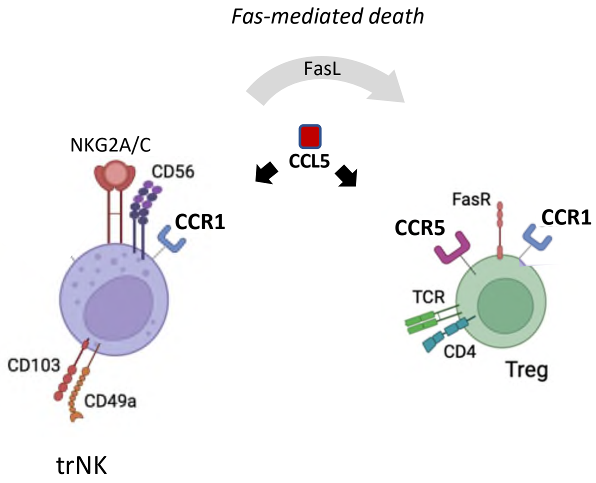

Figure supplement 8

A

Overall Survival

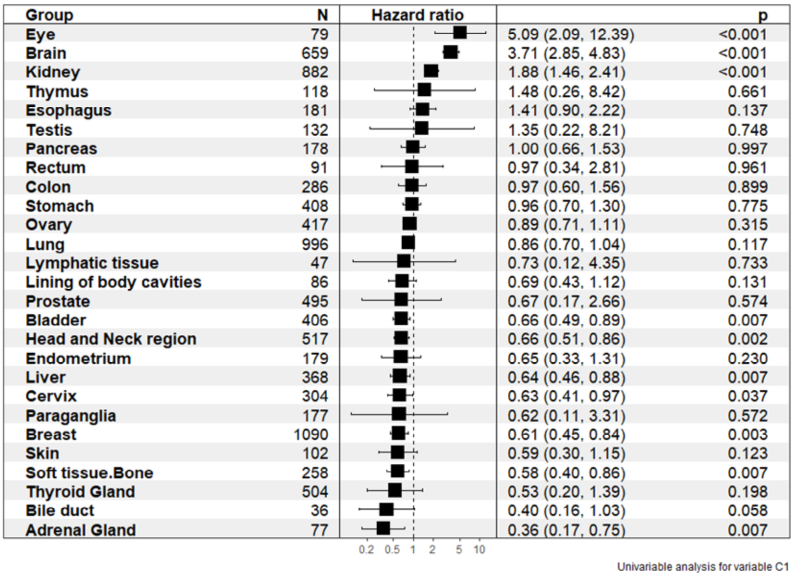


NK C1 – continuous variable

Disease Free Survival

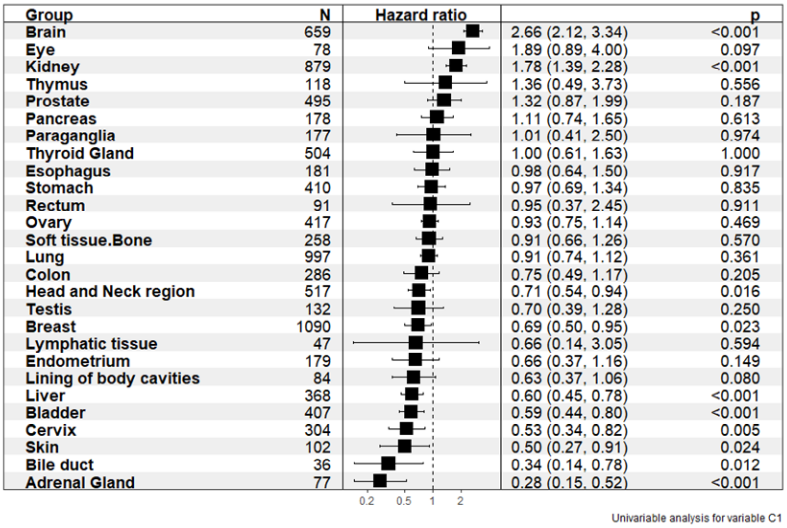


B

Overall Survival (univariate COX)

NK C1 (low v high)

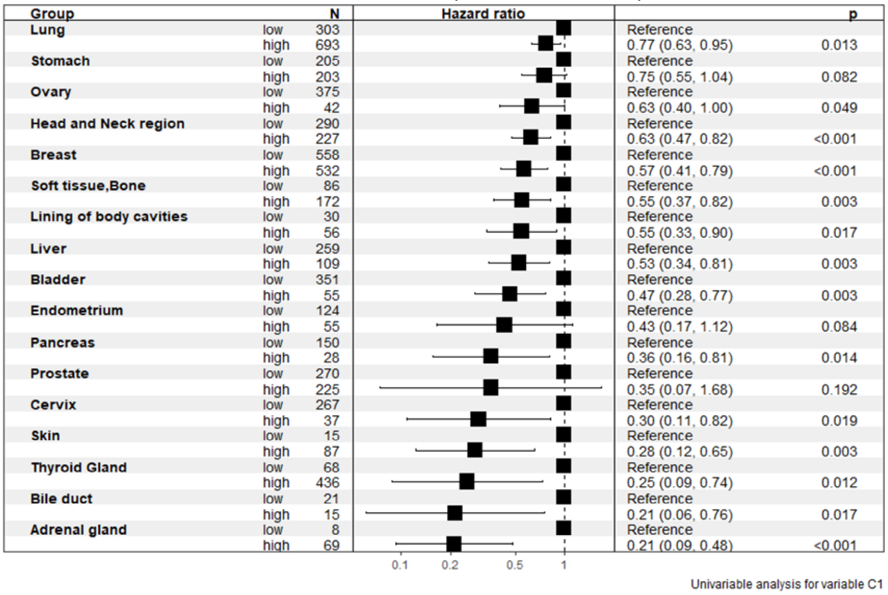

### Table supplement 1

NK_C1 vs ALL signature

| Genes |
|---|
| GZMA |
| TNFRSF18 |
| CD7 |
| CD247 |
| CTSW |
| HOPX |
| IL2RB |
| AC092580.4 |
| TMIGD2 |
| KLRB1 |
| NKG7 |
| CD96 |
| PTPN22 |
| LCK |

cDC1 vs ALL signature

| Genes |
|---|
| IDO1 |
| DNASE1L3 |
| CLEC9A |
| CLNK |
| ASB2 |
| FLT3 |
| P2RY14 |
| HLA-DOB |
| SERPINF2 |
| XCR1 |
| ENPP1 |
| CCSER1 |
| ZNF366 |
| GCSAM |
| TLR10 |
| BTLA |
| P2RY6 |
| PPM1J |
| MCOLN2 |

### Table supplement 2

Aurora Spectral Panel

| Antigen | Clone | Fluorochrome | Company, catalogue number | Dilution |
|---|---|---|---|---|
| Live/Dead | - | Blue | Thermofisher, L23105 | 1:1000 |
| CD45 | 30-F11 | AF532 | Thermofisher, 58-0451-80 | 1:50 |
| CD19 | 1D3 | SB436 | Thermofisher, 62-0193-80 | 1:50 |
| CD3 | 17A2 | eF450 | Biolegend, 48-0032-80 | 1:50 |
| NK1.1 | PK136 | BV650 | Biolegend, 108735 | 1:25 |
| CD8a | 53-6.7 | BV570 | Biolegend, 100739 | 1:25 |
| CD4 | RM4-5 | BV510 | BD Biosciences, 100553 | 1:100 |
| CD25 | 7D4 | AF647 | BD Biosciences, 563598 | 1:100 |
| FOXP3 | FJK-16s | PE | Thermofisher, 12-5773-80 | 1:20 |
| CD206 | C068C2 | BV785 | BD Biosciences, 141729 | 1:25 |
| F4/80 | T45-2342 | BV480 | BD Biosciences, 565635 | 1:100 |
| CD80 | REA983 | APC | Miltenyi Biotech, 130-116-46 | 1:100 |
| CD279 (PD-1) | J43 | PE-Cy7 | Thermofisher, 25-9985-82 | 1:25 |
| CD195 (CCR5) | REA354 | VB-FITC | Miltenyi Biotech, 30-105-141 | 1:20 |
| CD11b | M1/70 | AF700 | Biolegend, 101222 | 1:100 |
| Ly6C | HK1.4 | PE-Dazzle | Biolegend, 128043 | 1:200 |
| Ly6G | 1A8 | BV421 | Biolegend, 127627 | 1:25 |

### Table supplement 3

Multiplex IF panel

| Antigen | Company, catalogue (dilution) | Coupled to |
|---|---|---|
| NKG2D | Abcam, ab203353 (1:600) | Opal 520 |
| CD161/NK1.1 | Abcam, ab234107 (1:20000) | Opal 620 |
| CD3 | Abcam, ab5690 (1:300) | Opal 480 |
| CD8 | Cell Signalling, 98941 (1:800) | Opal 570 |
| E-cadherin | Cell Signalling, 3195 (1:500) | Opal 780 |
