## Supplementary material for "Tissue-resident NK cells support survival in pancreatic cancer through promotion of cDC1-CD8T activity": Figure Supplement Legends and Methods

**Figure supplement 1.**

(A) Human Protein Atlas derived single cell RNA-seq data of immune cells and relative CCL5 receptor expression (B) Overall Kaplan-Meier survival curves of PDAC patients (derived from TCGA cohorts) correlated to single specific immune gene signatures with the most opposing correlations (as shown in Figure 1C). Colour depicts positive (green) or negative (red) correlations. (C) Normalized expression of *NCAM1*, encoding neural cell adhesion molecule 1 (more commonly known as CD56); and *NCAM2*, encoding neural cell adhesion molecule 2, in different cells from the brain and immune cells. Data are derived from the Human Protein Atlas derived single cell RNA sequencing dataset (left). Overall Kaplan-Meijer survival curve of PDAC) does not significantly correlate with NCAM2 expression (right).

**Figure supplement 2**

(A) Representative H&E staining of pancreatic tumors derived from orthotopically injected KPC-A, KPC-F and KPC-G cells 30 days post injection (left). Representative immunofluorescence of KPC_A, KPC_F and KPC_G in vitro cultured cells stained with DAPI (blue; nucleus) and stromal markers (Collagen I; red, αSMA; cyan). (B) Chemiluminescent Cytokine array for KPF_F cell line (C) Optimized panel for immune-profiling using multispectral flow cytometry (Aurora Cytek) and gating strategy of respective cells as shown in Figure 2B.

**Figure supplement 3**

(A) Representative images of cross-sectional anatomy of a mouse abdomen, obtained through respiratory gated MRI (left), SARRP (centre; labelled CT), and an overlay of the MRI and CT images for tumor-specific dosage of 4 Gy (right). (B) Gross tumor volume at day 30 post-treatment as measured by MRI of mice as treated in Figure 3A. Data are represented as mean tumor volume±SD. (C) Quantitative measurement of DAPI+ cells based on H&E staining of mice treated as in Figure 3A. (D) Percentage of αSMA+ cells based on IHC (E) Quantification of IHC slides showing the percentage of CD8+ T-cell infiltrating the tumor (left). Tumor borders are marked as dashed lines. T=tumor, NT = non-tumor (right). (F) Profiles of infiltrating immune cells in pancreatic tumors by all treatments including single control treatments as measured by Aurora Cytek. Single, live cells were included for analysis and represented as frequencies of CD45+ cells in percentage.

**Figure supplement 4**

(A) Representative image of a multiplex immunofluorescence staining of a pancreatic tumor (B) Quantification of CD8+ cells (identified as DAPI+CD3+CD8+) across the different treatment groups. Markers are as indicated. Significance was tested for p<0.05 with a two-way ANOVA, *p>0.05, ***p>0.005. (C) Percentages of total DAPI+ immune cells (left) and E-cadherin+ tumor cells (right) derived from Figure 4A following treatment. Significance was tested using two-way ANOVA with Tukey multiple comparison (blue significance lines), or one-way ANOVA with Tukey multiple comparison (yellow significance lines) using a p<0.05. (D) Percentage of NKG2D+ and NKG2D- cells as a proportion of NK1.1+ cells as derived from Figure 4A. (E) UMAP of intratumoral NK cells (NK1.1+CD3-CD19-) in mice treated with the IR/CCR5i+αPD1 combination. (F) Representative example of gating strategy for the identification of tissue-resident (CD103+CD49a+) and conventional (CD103-CD49a-) NK cells. (G) Proportion of trNK, cNK, CD103+ NK, and CD49a+ NK in mice treated with the IR+IT regimen and control (mock), as a fraction of total NK cells

**Figure supplement 5**

(A) UMAP of the main cell types from the integrated scRNA-seq Steele dataset (left) and dot plot showing canonical marker expression for each respective cell type (right). Colour indicates average normalized expression; dot size represents percentage of cells of each cell type expressing the respective gene. (B) UMAP plot of lymphocyte sub-clusters from the Steele dataset (left) and their marker expressions (dot plot, right). (C) Violin plots showing NK-related gene signatures for each NK sub-cluster. (D) UMAP plot of lymphocyte sub-clusters from the Peng dataset, with the arrows representing the proposed functional development. (E) UMAP showing the expression levels of CD16(*FCGR3a*), CD56 (*NCAM1*), CD103 (*ITGAE*), and CD49a (*ITGA1*) across the NK sub-clusters. (F) Dot plot showing the different gene expression programs across the NK sub-clusters from the Peng dataset.

**Figure supplement 6**

(A) UMAP of the myeloid cell types from the Steele dataset (top) and the dot plot showing the canonical cell markers (bottom). (B) Heatmap of the top 15 markers for each DC sub-cluster. (C) Dot plot showing the estimated outgoing secreted signals from the NK C1 cell population. (D) Dot plot showing the estimated secreted and cell-cell contact signals from the cDC1 cell population. Colour of the dots represent the communication probability, and the size of the dot represents the significance level.

**Figure supplement 7**

(A) trNK cell (NK_C1) signature and overall survival in PDAC using the CPTAC3 dataset (B) Schematic overview of proposed mode of action of CCL5 gradients: CCL5 leads to the recruitment of CCR5+ immune cells, including Treg cells. Blockade of the CCL5-CCR5 axis however by CCR5 antagonists, allows recruitment of trNK cells via CCL5-CCR1 signalling whilst Tregs recruited into the microenvironment are killed off via FasL-mediated trNK killing.

**Figure supplement 8**

(A) Cox regression of trNK cell signature versus overall survival and disease-free survival across indicated malignancies**.**

**Table supplement 1**

The genetic signature of tissue-resident NK cells derived from scRNA-seq data of Figure 5C.

**Table supplement 2**

Antibody list of optimized panel for multispectral flow cytometry

**Table supplement 3**

Antibody list of optimized panel for multiplex immunofluorescence

**Materials & Methods**

### Mice and cell culture

C57BL6 mice (6 to 8 weeks) were purchased from Charles River (Kent, UK) and maintained in accordance with the UK Animal Law (Scientific Procedures Act 1986). NKPCB-NBC12.1F (KPC-F) cells, derived from a KPC mouse (LSL-Kras^G12D^; LSL-Trp53^R172H^; Pdx1^Cre^) derived from a CD57BL6 background were cultured in DMEM (Gibco) supplemented with 10% FBS, 2 mM glutamine, 100 U/mL penicillin, 100 ug/mL streptomycin and maintained under 5% CO_2_, 37°C.

### Orthotopic surgery

A total of 500 KPC-F cells (passages 9-11) were injected into C57BL6 mice in a volume of 5 µL in a mixture of 94% Matrigel and 6% DMEM (Gibco) using a 26-gauge syringe (Hamilton). Following anaesthesia of the mice using 4% isoflurane/oxygen and clipping of hair, an incision of 2cm was made in the left abdominal site. The spleen and pancreas were exteriorized using forceps and KPC-F cells were injected in the tail of the pancreas. Successful injection was verified by the appearance of a wheal at the injection site with no leakage through the pancreatic capsule. Spleen and pancreas were gently moved back into the peritoneal cavity; the peritoneum was sutured with absorbable 4.0 Vicryl suture (Ethicon) and the skin was closed using a 7 mm wound clip.

### *In vivo* drug treatments and radiation

Animals were randomized to different treatment groups following orthotopic implementation. For radiation, mice were anaesthetized and locally irradiated with 4 Gy using a combination of MRI and the small animal radiation research platform (SARPP). The CCR5 inhibitor Maraviroc (R&D) was administered intraperitoneally (10 mg/kg in PBS) from day 0 until day 6. Monoclonal Ultra-LEAF™ purified anti-PD1 antibody (BioLegend) was administered (5 mg/kg in PBS) via IP injection on alternate days from day 1 until day 7. Mice were culled at 30 days or if maximum tumor volume was reached.

### Monitoring of tumor growth via MRI

Mice were anesthetized using 1.5%-3% Isoflurane, positioned into a custom-made, 3D printed, multimodality cradle and scanned using a 4.7 T 310 mm horizontal bore Biospec AVANCE III HD preclinical imaging system equipped with 114 nm bore gradient insert (Bruker BioSpin GmbH, Germany). Respiration was maintained at 40-60 breaths/minute and monitored using a pneumatic balloon (VX010, Viomedex Ltd, UK) coupled to a pressure transducer and placed against the chest. A threshold-based respiration gating control signal was generated on a custom-built gating device, which allows efficient multi-slice MRI scanning with limited interference of respiratory motion^1^. Tumor volumes were calculated using ITK-SNAP^2^.

### Blood and tissue collection

Following reaching (humane) endpoints, mice were euthanized by approved Schedule 1 methods. Blood (≤1 mL) was collected via cardiac puncture under anaesthesia (3-3.5% Isoflurance/Oxygen) using a 25-gauge syringe (Thermo Fisher Scientific) coated with anticoagulant ACD buffer (0.5M glucose, 0.5M trisodium citrate, 0.5M citric acid; 100 µL). A total of 700 µL of blood was lysed with 12 mL of 1X RBC Lysis buffer and incubated for 10-15 minutes on a rotor, centrifuged, and washed with buffer (2% FBS, 1 mM EDTA in PBS). Three transverse sections of orthotopic tumors were taken and used for flow cytometry or fixed and stored in 10% neutral buffered formalin. Tumors were disassociated using a mixture of collagenase I, II and IV (Worthington) and DNAse (Thermo Fisher Scientific) in HBSS (Gibco), and continuously mixed at 850rpm for 45 minutes at 37°C, followed by rigours pipetting and additional dissociation for 50 minutes at 850rpm and 37°C. The disassociated tissue was strained through a 70 µm cell strainer and neutralized with DMEM+10% FBS prior to incubation with flow antibodies.

### Multicolor spectral flow cytometry

A total of 1x10^6^ cells for blood and disassociated tissue were surface stained in a 100 µL volume at 4°C in the dark (see Table 2 for list of antibodies and optimized dilutions). Cells and reference controls were centrifuged at 500*xg* for 2 minutes at room temperature and fixed and permeabilized using the FOXP3 Fixation/Permeabilization kit (Thermo Fisher Scientific). Samples were intracellularly stained at 4°C in the dark, washed, re-suspended in buffer and stored at 4°C in the dark prior to acquisition. Samples were acquired on a 4-laser Aurora (Cytek) with SpectroFlo software (Cytek, v2). A minimum of 10,000 events for reference control samples and 100,000 of CD45-gated events were recorded. Samples were acquired on a 3-laser (42-channel, V-16, B-16, R-10) or 4-laser (58-channel, UV-16, V-16, B-16, R-10) Cytek Aurora.

### Immunohistochemistry

A total of 24 hours post fixation at room temperature, tissues were transferred to 70% ethanol and stored overnight at 4°C. Fixed tissues were overnight processed using the STP120 Spin Tissue Processor (Thermo Fisher Scientific) and embedded the next day in paraffin wax. Tissues were cut into 4 µm sections using a Leica RM215 microtome and adhered onto SuperFrost Ultra Plus Adhesion slides (Thermo Fisher Scientific) before being dried overnight in a 37°C incubator.

### Multiplex immunofluorescence

Multiplex immunofluorescence staining was carried out on 4 µm thick FFPE sections by the Translation Histopathology Laboratory (THL, University of Oxford). In brief, sections were stained using the OPAL™ protocol (AKYOA Biosciences) on a Leica BOND RXm Auto-Stainer (Leica, Microsystems). Six consecutive staining cycles were performed using primary antibody-Opal fluorophore pairings detailed in Table 3. Primary antibodies were incubated for 30 minutes and detected using the BOND™ Polymer (Lecia Biosystems) as per manufacturer’s instructions. In brief, sections were baked, dewaxed with BOND™ dewax solution, rehydrated with alcohol and incubated with Epitope Retrieval Solution 1 or 2 (ER1, ER2) (Lecia Biosystems) at 100°C for 20 minutes. Sections were washed X3 with BOND™ wash, blocked with peroxidase block (3 – 4% (v/v) hydrogen peroxide) for 5 minutes and subsequently washed 3X with BOND™ wash. Primary antibodies were incubated for 30 minutes, washed as before, and incubated with Anti-Rabbit Poly-HRP IgG for 8 minutes. Sections were washed twice in BOND™ wash, once in deionized water prior to Opal antibody incubation for 10-minutes. Sections were washed three times in deionized water and finally incubated with spectral DAPI (Akoya Biosciences) and slides mounted with VECTASHIELD® Vibrance™ Antifade Mounting Medium (Vector Laboratories). Whole slide multispectral images were obtained on the AKOYA Bioscience Vectra® Polaris™ (scanned at 20X magnification). Batch analysis and spectral unmixing of the tissues was performed with inForm 2.4.11 software. Batched analysed multispectral images were fused in HALO AI to produce a spectrally unmixed reconstructed whole tissue image.

### Data analysis and statistics

Flow cytometry data was analysed using FlowJo software. HALO AI Software was used for analysis of multiplex immunofluorescence. Due to variations in staining, thresholds for individual colors were manually determined for individual tissue sections.

Bulk RNA-seq analysis

Gene expression, and clinical data were obtained via the TCGAbiolinks (2.24.3) R package for TCGA-PAAD^3^, and GDC data portal (<https://portal.gdc.cancer.gov/repository>) for CPTAC-3^4^. Raw read counts were transformed to counts-per-million (CPM) and log-CPM using the ‘edgeR’ Bioconductor package (3.38.0). Inter-sample variation was normalised using the trimmed mean of M-values (TMM) method of ‘edgeR’. The ‘GSVA’ package (1.44.0) was used to generate the tissue-resident NK signature (Table 1) score for individual patients. Gene signature scores and gene expressions were transformed to categorical groups via the maximally selected rank statistics (maxstat) method of the ‘survival’ package. Kaplan–Meier plots were performed using the ‘survival’ (3.3-1), and ‘survminer’ (0.4.9) R packages and the log-rank test was used to statistically compare survival estimates between groups.

TCGA Pan-cancer analysis

For the pan-cancer TCGA survival analysis, gene expression and clinical data were obtained via the UCSC Xena platform (<https://xenabrowser.net/datapages/?cohort=TCGA%20Pan-Cancer%20(PANCAN)&addHub=https%3A%2F%2Ftreehouse.xenahubs.net&removeHub=https%3A%2F%2Fpcawg.xenahubs.net>). The tissue-resident NK signature was analysed for the individual cancer types for Kaplan-Meier survival similar to above. Univariate COX regression of the NK signature for both overall survival and disease-free survival was performed using the `ezcox` (1.0.2) package.

Analysis of human single cell RNA-seq data

Raw scRNA-seq data from the Steele^5^ dataset were obtained from the NIH GEO database by using the accession number GSE155698. The samples included 16 treatment naïve primary tumors (6 surgical resections, and 10 fine-needle biopsy specimens) and three non-malignant pancreas samples. All analysis was performed using the `Seurat` (4.3.0) package. Cells with low number of detected genes (<200), high number of mitochondrial genes (>25%), and high number of gene counts (>100,000) were filtered out from further analysis. Data were integrated to remove batch effects using a reciprocal principal component analysis (rPCA) based on the developer’s pipeline (<https://satijalab.org/seurat/articles/integration_rpca.html>). Following integration, data were scaled, and visualized using uniform manifold approximation and projection (UMAP) on the top 40 principal components. Cell clusters were identified using the `FindNeighbors` and `FindClusters` Seurat functions and annotated based on canonical cell markers. Gene expression differences across cell clusters were visualized using the `DotPlot` and `DoHeatmap` Seurat functions. Curated gene sets were calculated using the `AddModuleScore` and compared using violin plots with the `VlnPlot` function.

Inference of cell-cell interactions

The `CellChat`^6^ (1.6.1) R package was used to identify and visualize significant cell-cell interactions based on curated ligand-receptor pairs for both Secreted Signalling and Cell-Cell Contact categories. Network centrality scores were calculated with the `netAnalysis_computeCentrality` function and global communication patterns were visualized using the `netAnalysis_signalingRole_heatmap`. Specific interactions of cells of interest were visualized by circle and dot plots by using the `netVisual_circle` and `netVisual_bubble` functions, respectively.

Packages/tools used for analysis:

| Package (version) | Repository | Link to package |
| --- | --- | --- |
| TCGABiolinks 2.24.3 | Bioconductor | <https://academic.oup.com/nar/article/44/8/>  e71/2465925?login=true |
| edgeR 3.38.0 | Bioconductor | <https://academic.oup.com/bioinformatics/article>/26/1/139/182458?login=true |
| GSVA 1.44.0 | Bioconductor | https://bmcbioinformatics.biomedcentral.com/articles/10.1186/1471-2105-14-7 |
| Survival 3.3-1 | CRAN | <https://link.springer.com/book/10.1007/978-1-4757-3294-8> |
| Survminer 0.4.9 | CRAN | <https://rpkgs.datanovia.com/survminer/index.html> |
| Ezcox 1.0.2 | CRAN | https://github.com/ShixiangWang/ezcox |
| Seurat 4.3.0 | CRAN | <https://github.com/satijalab/seurat> |
| CellChat 1.6.1 | Github | <https://www.nature.com/articles/s41467-021-21246-9> |
| Tidyverse 2.0.0 | CRAN | <https://www.tidyverse.org/> |

Data availability statement
The authors confirm that the data supporting the findings of this study are available within the article, its supplementary materials or upon request. Sources of publicly available data and analysis packages are referenced with the text or methods.

Supplementary References

1. Kinchesh P, Allen PD, Gilchrist S, Kersemans V, Lanfredini S, Thapa A, O'Neill E, Smart SC**.** [Reduced respiratory motion artefact in constant TR multi-slice MRI of the mouse.](https://pubmed.ncbi.nlm.nih.gov/30928386/) Magn Reson Imaging. 2019 Jul;60:1-6.
2. Yushkevich PA, Yang Gao, Gerig G.TK-SNAP: An interactive tool for semi-automatic segmentation of multi-modality biomedical images. Annu Int Conf IEEE Eng Med Biol Soc. 2016:3342-3345
3. Cancer Genome Atlas Research Network. Integrated Genomic Characterization of Pancreatic

Ductal Adenocarcinoma. Cancer Cell 2017;32(2):185,203.e13.

4. Cao L, Huang C, Cui Zhou D, Hu Y, Lih TM, Savage SR et al. Proteogenomic characterization of

pancreatic ductal adenocarcinoma. Cell 2021;184(19):5031,5052.e26.

5. Steele NG, Carpenter ES, Kemp SB, Sirihorachai VR, The S, Delrosario L et al. Multimodal mapping

of the tumor and peripheral blood immune landscape in human pancreatic cancer. Nat Cancer 2020;1(11):1097-112.

6.     Jin S, Guerrero-Juarez CF, Zhang L, Chang I, Ramos R, Kuan C et al. Inference and analysis of cell-

cell communication using CellChat. Nat Commun 2021;12(1):1-20.
